# Component A2 is a redox-sensitive archaeal ATPase activated by methyl-coenzyme M reductase

**DOI:** 10.64898/2026.03.18.712670

**Authors:** Sophia A. Adler, Dipti D. Nayak

## Abstract

Methyl-coenzyme M reductase (MCR) is the primary source of biogenic methane on Earth. In the active site of MCR, a nickel (Ni)-containing porphyrin (F430) must be in the Ni^1+^ oxidation state to initiate catalysis. The reductive activation of MCR, i.e., reduction of F430 to its Ni^1+^ state, is an ATP-dependent process, but the underlying ATPase and its precise role remain unknown. Component A2 is an ATP-binding protein that associates with MCR but, since it was reported to lack ATPase activity, its putative function was designated as an ATP-carrier protein. In contrast, recent structural insights into the MCR activation complex suggest that component A2 might hydrolyze ATP to drive conformational changes required for enzyme activation. Here, we provide direct biochemical evidence that component A2 is a bona fide ATPase that hydrolyzes ATP under strictly anaerobic conditions and only upon interaction with MCR. Mutational analyses reveal that component A2 must be bound to ATP prior to association with MCR and that residues involved in ATP hydrolysis do not impact protein-protein interaction. The two nucleotide-binding domains of A2 act cooperatively but display asymmetric contributions to ATP hydrolysis and MCR engagement. In addition, a distinctive N-terminal zinc-binding motif (ZBM) is required for maximal ATPase activity but is dispensable for MCR binding. Phylogenetic analyses reveal that this ZBM distinguishes component A2 from related ABC-type ATPases. Together, these findings identify component A2 as a distinct class of remodeling ATPases that powers conformational changes underlying the reductive activation of MCR.

**Significance Statement:** A large fraction of methane on Earth is generated by methanogenic archaea using the enzyme methyl coenzyme-M reductase (MCR). The maturation of MCR is a multi-step ATP-dependent process but the role of ATP and the corresponding ATPase(s) have remained unclear. Here, we show that component A2, a protein that is universally conserved in archaea that encode MCR and related enzymes, hydrolyzes ATP only upon interaction with MCR under anaerobic conditions. Our findings, together with recent structural studies, indicate that component A2-mediated ATP hydrolysis facilitates the reductive activation of MCR during the final step of its maturation. These results clarify a key step in the biogenesis of a central enzyme involved in biological methane production.

## Introduction

Methyl-coenzyme M reductase (MCR)-encoding archaea plays a prominent role in the global methane cycle by catalyzing a reversible reaction shown below (1):

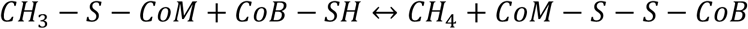

In methanogenic archaea, MCR mediates the final step of methanogenesis to generate methane. In anaerobic methanotrophic archaea (ANME), MCR activates methane to methyl-coenzyme M, which is the first step in its eventual oxidation to carbon dioxide (2, 3). Despite substantial sequence-level divergence, the overall structure and active site architecture of MCR is extremely well-conserved across archaea (4). The ∼300 kDa MCR heterohexamer is comprised of three subunits arranged in an α_2_β_2_γ_2_ conformation and contains two active sites that are 50 Å apart and proposed to be functionally coupled (Fig. 1) (4–6). Each active site has a non-covalently bound prosthetic group, factor 430 (F430), that is only found in MCR and other members of the alkyl-coenzyme M reductase (ACR) enzyme family (2, 4, 7, 8). F430 is a Ni-containing tetrapyrrole that must be reduced to its Ni^1+^ oxidation state, through a process known as reductive activation, for MCR to be catalytically active (9–11). As the mid-point potential of the Ni^2+^/Ni^1+^ couple is extremely low (between -600 mV and -700 mV relative to a standard hydrogen electrode) (12), the reductive activation of MCR requires many accessory proteins (Fig. 1) (13, 14). In the absence of these proteins, the Ni atom in F430 gets rapidly oxidized to Ni^2+^ and MCR becomes catalytically inactive, even when purified under anaerobic conditions (15).

**Figure 1.**
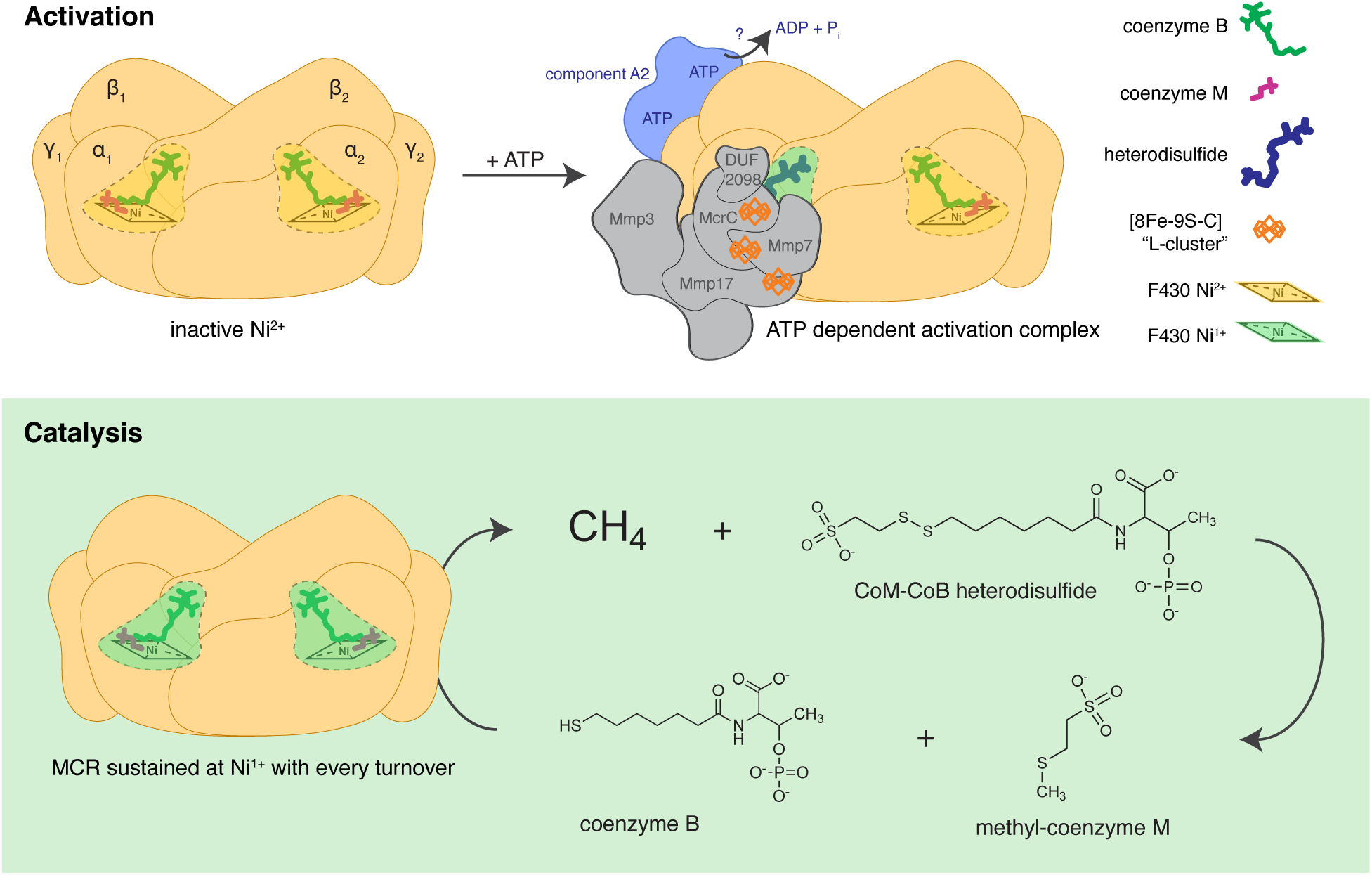
Cartoon depicting the reductive activation and catalytic activity of MCR. (Top) Reductive activation of MCR is an ATP-dependent process. Component A2 (light blue) and other methanogenesis marker proteins or MMPs (gray) are required to generate the Ni^1+^ form of F430 (green) for activation of MCR from its Ni^2+^ oxidation state (yellow) to its Ni^+1^ oxidation state (green). (Bottom) The reaction catalyzed by MCR requires an external input of electrons for the reduction of the methyl-group in methyl-coenzyme M to methane, which is provided by coenzyme B. The Ni^1+^ form of F430 is regenerated at the end of each reaction cycle so ATP-dependent activation is not required for each molecule of methane generated.

For nearly fifty years it has been known that the reductive activation of MCR is an ATP-dependent process, but the underlying reason is not clearly understood. In 1978, Gunsalus and colleagues showed that *Methanothermobacter thermoautotrophicus* cell extracts required trace amounts of ATP for methane production from methyl-coenzyme M and estimated that *∼*1 mol of ATP was required for ∼15 mols of methane produced (16). Subsequently, Rouvier et al. isolated an ATP-binding protein, designated as component A2 or AtwA, that was required for optimal methane production by cell extracts of *M. thermoautotrophicus* (17). Initial reports indicated that component A2 is a soluble, colorless, air-tolerant protein of the ABC-type ATPase family that lacks any detectable ATP hydrolysis activity (17, 18). Component A2 could not perform electron transfer either to or from common electron carriers like FAD, NAD, NADP, or F_420_ (17). Based on these analyses, component A2 was designated as an ATP-carrier protein that delivers ATP to another enzyme, often speculated to be a homolog of the dinitrogenase reductase (NifH) (19), which can hydrolyze ATP and transfer electrons to F430 for the reductive activation of MCR. However, NifH homologs have not, to our knowledge, been shown to copurify with MCR even though they are universally conserved in MCR-encoding archaea (20, 21).

Recently, a cryo-EM structure of the MCR activation complex shed new light on the putative function of component A2 (13). The MCR activation complex contains component A2 bound asymmetrically to one γ subunit (McrG) (Fig. 1). In addition, the MCR activation complex also contains McrC and three other methanogenesis marker proteins (MMPs) – Mmp3, Mmp7, and Mmp17 – that coordinate three complex Fe-S clusters, which closely resemble the [8Fe-9S-C] L clusters involved in the maturation of nitrogenases (Fig. 1). Even though component A2 does not interact directly with McrC and the MMPs in the activation complex, it is hypothesized to stabilize the N-terminal “latch” of the α subunit (McrA) to make the active site more accessible for electron transfer to F430 through the putative L clusters. To this end, Ramirez-Amador and colleagues showed that addition of ATP stabilizes the MCR activation complex and postulated that this process is mediated by structural rearrangements that occur when component A2 hydrolyzes ATP. This hypothesis directly contradicts previous biochemical evidence that component A2 is merely an ATP-carrier protein. Furthermore, since component A2 is bound to ATP —not ADP — in the activation complex, it is still plausible that it delivers ATP to another ATPase domain protein, like MMP15, that could transiently associate with the activation complex, making it difficult to capture structurally (22).

Here, we resolve a longstanding conundrum regarding the role of component A2 in the reductive activation of MCR by complementary in vivo and in vitro analysis of the protein from *Methanosarcina acetivorans.* Our results show that component A2 is a bona fide ATPase that hydrolyzes ATP only upon interaction with MCR under anaerobic conditions, and that interaction with MCR is predicated on component A2 first binding ATP. In addition, our evolutionary analyses indicate that component A2 belongs to a unique class of redox-active remodeling ATPases that have co-evolved with members of the ACR family within archaea.

## Results

### Component A2 is essential and constitutively expressed in *Methanosarcina acetivorans*

In *M. acetivorans*, component A2 is encoded by MA_3998 (MA_RS20860) and is found in the vicinity of six genes encoding methanogenesis marker proteins (MMPs) (Fig. 2a), some of which have also been implicated in the activation of MCR (Fig. 2b), and all of which are highly conserved across genomes containing MCR/ACR (13, 23). Component A2 and these six others neighboring MMPs are often referred to as the MCR activation operon, but it is unclear if these genes are co-transcribed and the degree to which they are expressed. First, to test if component A2 is in an operon with the other six MMPs (shown in Fig. 2a), we extracted RNA from *M. acetivorans* and performed RT-PCR with primers that span intergenic regions across the putative operon as shown in *SI Appendix*, Fig. S1a. We were able to observe PCR products for all the intergenic regions tested, which experimentally validates that these seven genes are, indeed, organized as an MCR activation operon (*SI Appendix*, Fig. S1a). Next, we mapped RNA-sequencing reads from previous transcriptomics experiments with *M. acetivorans* (24–26) to the MCR activation operon. The average expression of the MCR activation operon does not vary significantly across growth substrates and remains unchanged in response to MCR-limitation (*SI Appendix*, Figs. S1b and S1c). The read depth and coverage of the MCR activation operon is ∼40-fold lower than the MCR operon (*mcrBDCGA*) (Fig. 2c and *SI Appendix*, Fig. S2). Taken together, these data suggest that component A2 is part of the MCR activation operon, which is constitutively expressed at much lower levels than MCR operon in *M. acetivorans*.

**Figure 2.**
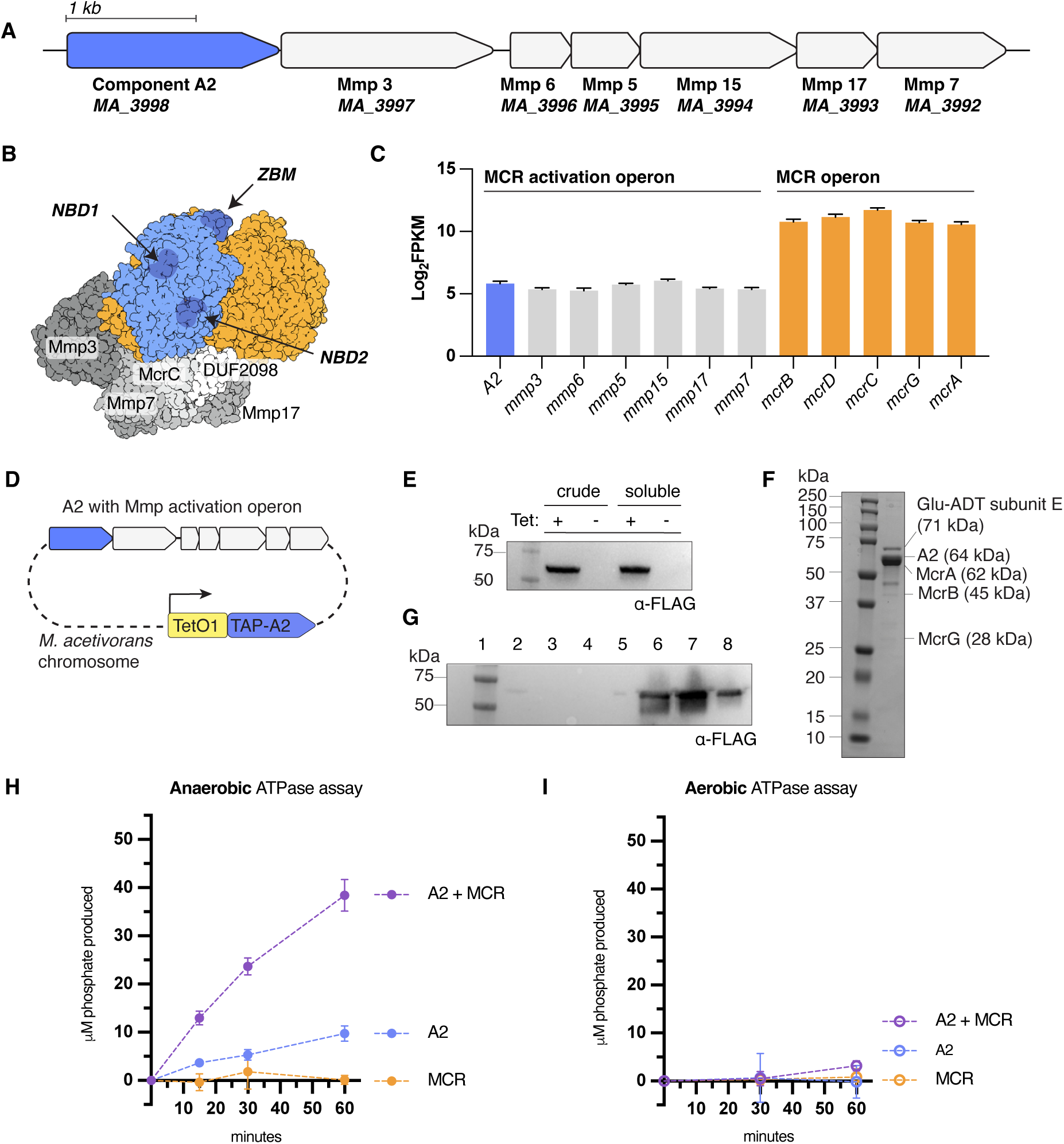
Component A2 is an oxygen-sensitive ATPase that hydrolyzes ATP upon interaction with MCR. (a) Genomic arrangement of the MCR activation operon in *M. acetivorans* with scale bar depicting 1 kilobase pair. (b) Structure of the putative MCR activation complex (PDB: 9H1L) with MCR in orange, component A2 (with relevant domains highlighted) in blue, and the other methanogenesis marker proteins in gray. NBD refers to **N**ucleotide **B**inding **D**omain and ZBM refers to **Z**inc **B**inding **M**otif. (c) Log_2_ transformed FPKM (**F**ragments **P**er **K**ilobase of transcript per **M**illion mapped reads) of the MCR activation operon and the MCR operon in *M. acetivorans* grown in high-salt (HS) minimal medium supplemented with trimethylamine (TMA) at 37 °C. (d) Genotype of an *M. acetivorans* strain expressing a second copy of component A2 in trans under the control of a tetracycline inducible promotor. (e) Anti-FLAG immunoblot showing the inducible production of component A2 upon the addition of 100 µg/mL tetracycline (tet) to the growth medium in crude and soluble cell lysates with 13.9 µg total protein loaded into each lane. (f) Anti-FLAG immunoblot of aerobic affinity-purification of A2 with a Streptactin resin. The lanes represent the following: (1) ladder, (2) cell lysate, (3) flow-through, (4) first wash, (5) second wash, (6) first elution, (7) second elution, and (8) third elution. (g) SDS-PAGE gel showing anaerobic purification of full-length TAP-tagged component A2 (64 kDa with tag). (h) Anaerobic ATPase assay of component A2 alone (blue), MCR alone (purple), and component A2 combined with MCR (orange). Each reaction contained 500 µg/mL of each protein indicated with 200 µM ATP, 10 mM MgCl_2_, 20 mM HEPES, 300 mM NaCl, and 1% glycerol. Reactions were incubated at 37 °C. Inorganic phosphate production was measured at 0, 15, 30, and 60 minutes using the malachite green reagent. (I) Aerobic ATPase assay of component A2 alone (blue), MCR alone (purple), and component A2 combined with MCR (orange). Proteins were purified anaerobically then removed from the anaerobic chamber and reactions were set up on the bench top. The same assay conditions as (c) were used, but time points were taken at 0, 30, and 60 minutes. Error bars represent the standard deviation of three technical replicates for each reaction.

Based on their proposed function, component A2 and other genes in the MCR activation operon are likely to be essential in methanogens, but this hypothesis, to the best of our knowledge, has not been explicitly tested. To test the essentiality of the MCR activation operon, we designed CRISPR-editing plasmids to generate in-frame chromosomal deletions of each gene individually (27). For component A2, transformation of the CRISPR-editing plasmids did not produce any viable colonies (*SI Appendix*, Fig. S3). For the rest of the genes- Mmp5, Mmp6, Mmp17, Mmp7, Mmp3, and Mmp15- in the MCR activation operon, a few antibiotic-resistant transformants were detected (*SI Appendix*, Fig. S3). However, PCR with primers flanking the gene of interest revealed either a mixed population of the wildtype (WT) and the mutant allele or only the WT allele, suggesting that genome editing was either incomplete or unsuccessful, respectively (*SI Appendix*, Fig. S3). A similar observation was also made in recent attempts to delete *nifB* (28), another essential gene in *M. acetivorans*. Altogether, we were unable to successfully delete component A2 or any of the neighboring MMPs in *M. acetivorans,* which corroborates their proposed role in the activation of MCR, an enzyme essential for growth and viability.

### Component A2 hydrolyzes ATP after interacting with MCR under anaerobic conditions

Since the native component A2 locus is essential and constitutively expressed at low levels in *M. acetivorans* (Fig. 2c), we generated a strain to overexpress it in trans for facile purification and biochemical characterization of its proposed function (Fig. 2d). Expressing component A2 in trans also allowed us to study catalytically inactive mutants as the chromosomal locus retained its original function, which, as noted above, appears to be essential for cell viability. To this end, we fused a **<u>t</u>**andem-**<u>a</u>**ffinity **<u>p</u>**urification (TAP) tag, comprised of a Twin-Strep-1X-FLAG sequence, at the N-terminus of component A2 and expressed it under the control of the P*mcrB*(tetO1) tetracycline inducible promoter in the pJK027A vector backbone (29). We chose to add a TAP tag the N-terminus of component A2 as it is far from the interaction interface with MCR and is also known to not disrupt ATP-binding (13, 19). The resulting plasmid (pSAA004) was integrated in the chromosome of *M. acetivorans* (WWM73) at the **φ**C31 attachment site as described previously (29). Whole genome sequencing was performed to verify the genotype of the strain used for protein purification (*SI Appendix*, Table S1) and the production of TAP-tagged component A2 was detected by immunoblotting against the FLAG-tag when expression was induced with 100 µg/mL tetracycline (Fig. 2e). Overexpression of component A2 did not lead to an observable growth defect, which indicated that elevated levels of this protein are not toxic or lethal to *M. acetivorans* (*SI Appendix*, Fig. S4). Anaerobic affinity purification of TAP-tagged component A2 yielded colorless, soluble, full-length protein of *∼* 64 kDa (Figs. 2f, 2g and *SI Appendix*, Figs. S5a, S5b, S5d). In addition to component A2, we observed nine additional bands that either correspond to McrB, McrG or degradation products of component A2 itself (Fig. 2f and *SI Appendix*, Table S2). A band corresponding to Glutamyl-tRNA(Gln) amidotransferase subunit E (*SI Appendix*, Table S2) was observed in all protein preparations of component A2 (even the mutant forms; see below) hence was not considered to impact any of the data presented here. Since McrA is the same size as component A2, we used an anti-McrA antibody to confirm that it also copurifies with this protein (*SI Appendix*, Fig. S5e). A reverse pulldown with TAP-tagged MCR (*SI Appendix*, Fig. S5c) confirmed the interaction between component A2 and MCR (*SI Appendix*, Table S3).

While it is well established that component A2 binds ATP (13, 17, 30), it is unclear if its role in the context of MCR activation is to serve as an ATP-carrier for an alternate ATPase or to perform ATP hydrolysis by itself (19). To disentangle these two hypotheses, we explored if component A2 can hydrolyze ATP in isolation or in conjunction with MCR using the Malachite green assay to quantify free inorganic phosphate (Pi) produced during the process (see more details in the Materials and Methods section). When component A2 was purified under anaerobic conditions, we detected robust ATPase activity, ∼ 38 nmol Pi released/mg protein after 60 minutes of incubation for three independent preparations (Figs. 2h, 3j and *SI appendix*, Fig. S5f). In contrast, when we purified component A2 under aerobic conditions, no free Pi was detected under the same conditions (*SI appendix*, Fig. S5f). To rule out the possibility that the ATP hydrolysis in anaerobic preparations of component A2 was due to co-purifying contaminant(s), we assayed the ATPase activity of anaerobically prepared A2 after air exposure. Treatment with air and oxic buffers abolished ATPase activity (Fig. 2i), which suggests that component A2 is an oxygen-sensitive protein and can only hydrolyze ATP under reducing conditions.

**Figure 3.**
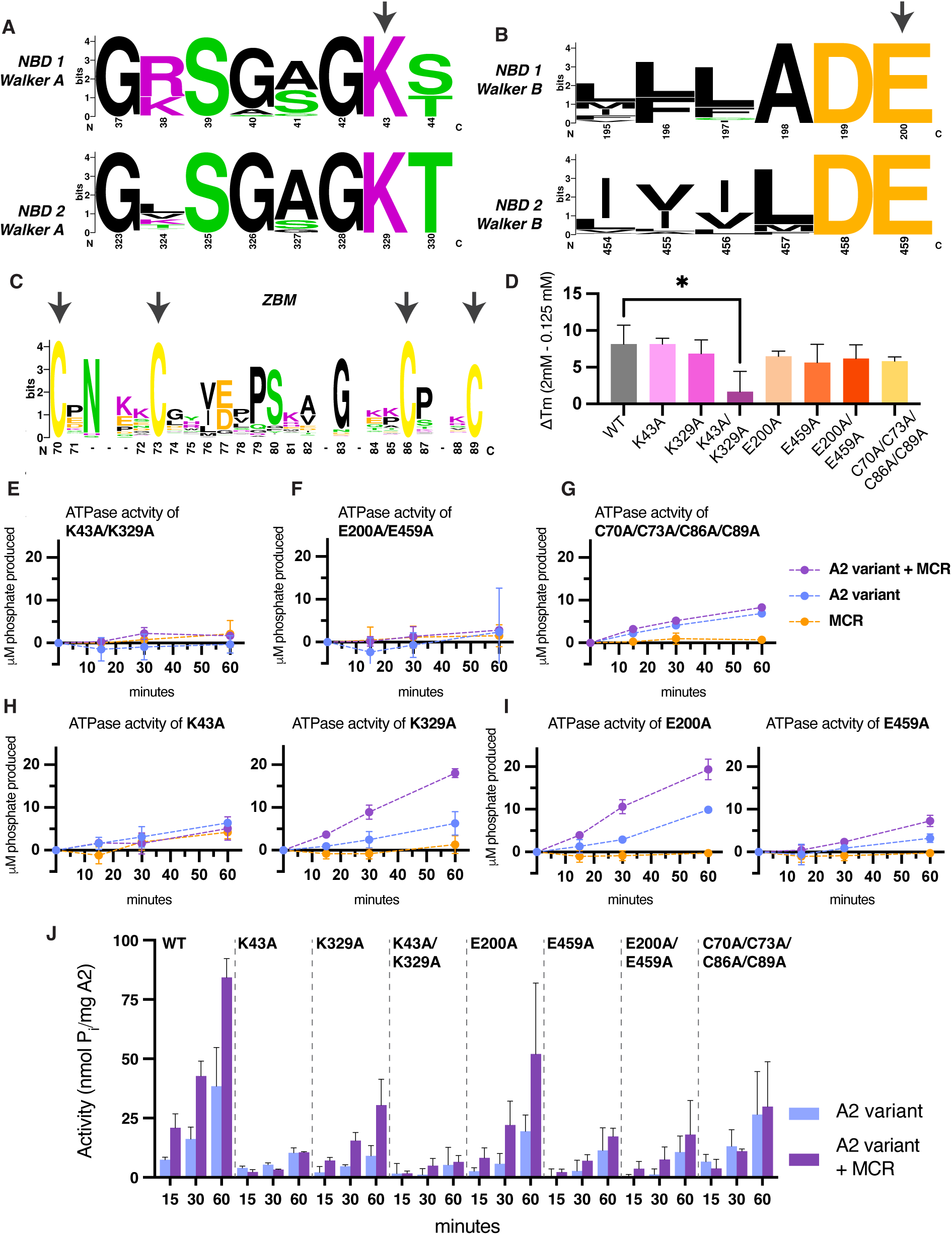
Nucleotide binding domains (NBDs) and zinc binding motif (ZBM) of component A2 are highly conserved and disrupt ATPase activity. (a-c) Sequence alignment of 888 component A2 sequences assigned to HMM (Hidden Markov Model) TIGR03269. The x-axis indicates position number in *Methanosarcina acetivorans*. Arrows indicate the (a) mutated residues K43 and K329, (b) D200 and D459, and (c) C70, C73, C86, and C89. (d) Change in melting temperature (ΔTm) of component A2 variants in buffer with 2 mM ATP versus 125 µM ATP. Error bars represent standard deviation of the mean ΔTm of three independent preparations of the protein (except for E200A and E459A, which represent the data for two and four protein preparations, respectively) and Welch’s unpaired t-test was performed to assess statistical significance; * indicates P-value < 0.05. (e-i) Anaerobic ATPase assay of each indicated component A2 variant (blue), MCR alone (orange), and the component A2 variant combined with MCR (purple). Each reaction contained 500 µg/mL of each protein with 200 µM ATP, 10 mM MgCl_2_, 20 mM HEPES, 300 mM NaCl, and 1% glycerol. Reactions were incubated at 37 °C. Inorganic phosphate production was measured at 0, 15, 30, and 60 minutes using malachite green reagent. Error bars represent the standard deviation of three technical replicates. (j) ATPase activity for each component A2 variant alone and upon addition of MCR. Error bars represent the standard deviation of three independent preparations of each variant of A2, except C70A/C73A/C86A/C89A for which only two independent preparations were performed.

Next, we tested if the addition of MCR impacts ATP hydrolysis by component A2 in vitro. Anaerobic preparations of MCR from stationary-phase cultures of *M. acetivorans* had no detectable ATPase activity whereas ATP hydrolysis was observed in MCR preparations from exponential-phase cultures, likely because component A2 copurifies with it (*SI Appendix*, Fig. S5h). Hence, we only used MCR derived from stationary-phase cultures to test its impact on ATPase activity of component A2 (Fig. 2h). The addition of an equal amount of MCR (by weight) to component A2 increased its ATPase activity under anaerobic conditions by 2.2-fold (Figs. 2h, 3j, and *SI appendix*, Fig. S5g). In contrast, no detectable ATPase activity was observed when MCR was added to component A2 under aerobic conditions (Fig. 2i). These data are consistent with the hypothesis that the ATPase activity of component A2 alone is due to the small amount of MCR that copurifies with it, which is further enhanced by the addition of MCR. Taken together, our in vitro analyses suggest that component A2 is a redox-sensitive ATPase that hydrolyzes ATP only after it interacts with MCR.

### At least one nucleotide-binding domain and the zinc-binding motif are important for ATP hydrolysis by component A2

Component A2 from *M. acetivorans* contains two nucleotide binding domains (NBDs) as well as a CXXCX_12_CXXC Zn^2+^-binding motif (ZBM) (Figs. 3a-3c). The two NBDs are highly conserved in component A2 sequences across archaea and each of them contains a Walker A motif (GxxGxGK[T/S]) and Walker B motif (hhhhDE) (Figs. 3a and 3b). The lysine residue (K) of the Walker A motif is known to be critical for ATP-binding (31), and the glutamate residue (E) of the Walker B motif deprotonates water in the first step of ATP hydrolysis and is essential for catalysis (32). To eliminate ATP-binding, we mutated the lysine residues in the two Walker A motifs to alanine by generating a K43A/K329A double point mutant of component A2. To specifically abolish ATPase activity, without impacting ATP-binding, we generated a E200A/E459A double point mutant of component A2. Additionally, we generated single point mutants, K43A, K329A, E200A, and E459A, to determine if catalytic activity is dependent on both NBDs and if the two NBDs act asymmetrically. All point mutants were expressed and purified from *M. acetivorans* using the strategy described for the WT allele in Fig. 2 (*SI Appendix*, Figs. S6-S8).

To validate the proposed role of each of these NBD residues, we performed differential scanning fluorimetry (DSF) (or thermal shift assays) in the presence of low (0.125 mM) or high (2 m) concentrations of ATP in the buffer (33). All DSF assays were performed aerobically with anaerobically purified enzyme, and the first differential of the melt curve had a single, major peak, which we infer corresponds to the melting temperature (‘Tm’) of component A2 (Fig. 3d and *SI Appendix*, Fig. S9). Switching from low to high ATP concentrations in the buffer increased the Tm (referred to as the ∆Tm) of all mutants of components A2 by > 5.0 °C, apart from the K43A/K329A double mutant (Fig. 3d and *SI Appendix*, Fig. S9). These data suggest that ATP-binding stabilizes component A2 and that the K43A/K329A double mutant, as expected, does not experience this shift in Tm because it cannot bind ATP. Like the NBDs, the CXXCX_12_CXXC ZBM is highly conserved across all component A2 sequences too (Fig. 3c), which suggests that it is functionally relevant. To test the role of the ZBM, we generated a C70A/C73A/C86A/C89A mutant. The ZBM mutant was stabilized by the presence of ATP in the buffer (Fig. 3d), which indicates that it can still bind ATP.

Next, we measured the anaerobic ATPase activity of all the point mutants either by themselves or in the presence of an equal amount of MCR (by weight). The K43A/K329A double mutant produced < 6 nmol Pi/mg protein after 60 minutes of incubation, which did not increase upon the addition of MCR (Figs. 3e, 3j and *SI Appendix*, Fig. S10a). ATP hydrolysis was similarly attenuated in the E200A/E459A mutant (Figs. 3f, 3j and *SI Appendix*, Fig. S10b). The C70A/C73A/C86A/C89A mutant had ∼50% activity of the WT by itself, which did not increase upon the addition of MCR (Figs. 3g, 3j and *SI Appendix*, Fig. S10c). These data suggest that the NBDs are essential whereas the ZBM is critical for optimal ATPase activity.

To test if the two NBDs work synergistically or independently, we assayed the ATPase activity of the single point mutants. While neither single mutant behaved like WT, the K329A mutant had *∼* 2.9 times higher activity than the K43A mutant in the presence of MCR (Figs. 3h, 3j and *SI Appendix*, Figs. S11a, S11b). Similarly, the E200A mutant had *∼* 3 times higher activity than the E459A mutant (Figs. 3i, 3j and *SI Appendix*, Fig. S11c, S11d). These data suggest that the NBDs have an asymmetric but synergistic contribution to ATP hydrolysis.

### Only ATP-bound component A2 can interact with MCR

There are two models for ATP hydrolysis by the component A2-MCR complex that are equally consistent with the activity data noted above. One is that component A2 binds ATP first, ATP-bound component A2 interacts with MCR, and then ATP hydrolysis occurs (Model 1 in Fig. 4a). Alternately, component A2 binds MCR first, ATP binds to the component A2-MCR complex, which is followed by hydrolysis (Model 2 in Fig. 4a). To distinguish between these two possibilities, we leveraged DSF to observe complex formation between component A2 and MCR either in the presence or absence of different adenine nucleotides. All DSF assays were performed aerobically with anaerobically purified enzyme, so that the interaction between component A2 and MCR could be decoupled from ATP hydrolysis. In the presence of ATP, the first differential of the melt curve for component A2 and MCR individually showed a single peak at 45.5 ± 1.3 °C and 67.2 ± 0.8 °C respectively, which corresponds to the Tm of these two proteins (Figs. 4b, 4c, and *SI Appendix*, Figs. S12a, S12b). When component A2 and MCR are combined in the presence of ATP, a third peak at 60.8 ± 0.6 °C appears in the melt curve (Fig. 4d, and *SI Appendix*, Fig. S12c). We interpret that this third peak corresponds to the Tm of the complex of component A2 and MCR. This third peak is absent when MCR and component A2 are combined in the absence of ATP or in the presence of ADP (Figs. 4d, 4e, and *SI Appendix*, Figs. S12d-f). All tested concentrations of ATP showed some presence of this component A2-MCR peak (Fig. 4e and *SI Appendix*, Fig. S12g). In contrast, a peak corresponding to the component A2-MCR complex was not detected regardless of the [ADP] (Fig. 4e and *SI Appendix*, Fig. S12h).

**Figure 4.**
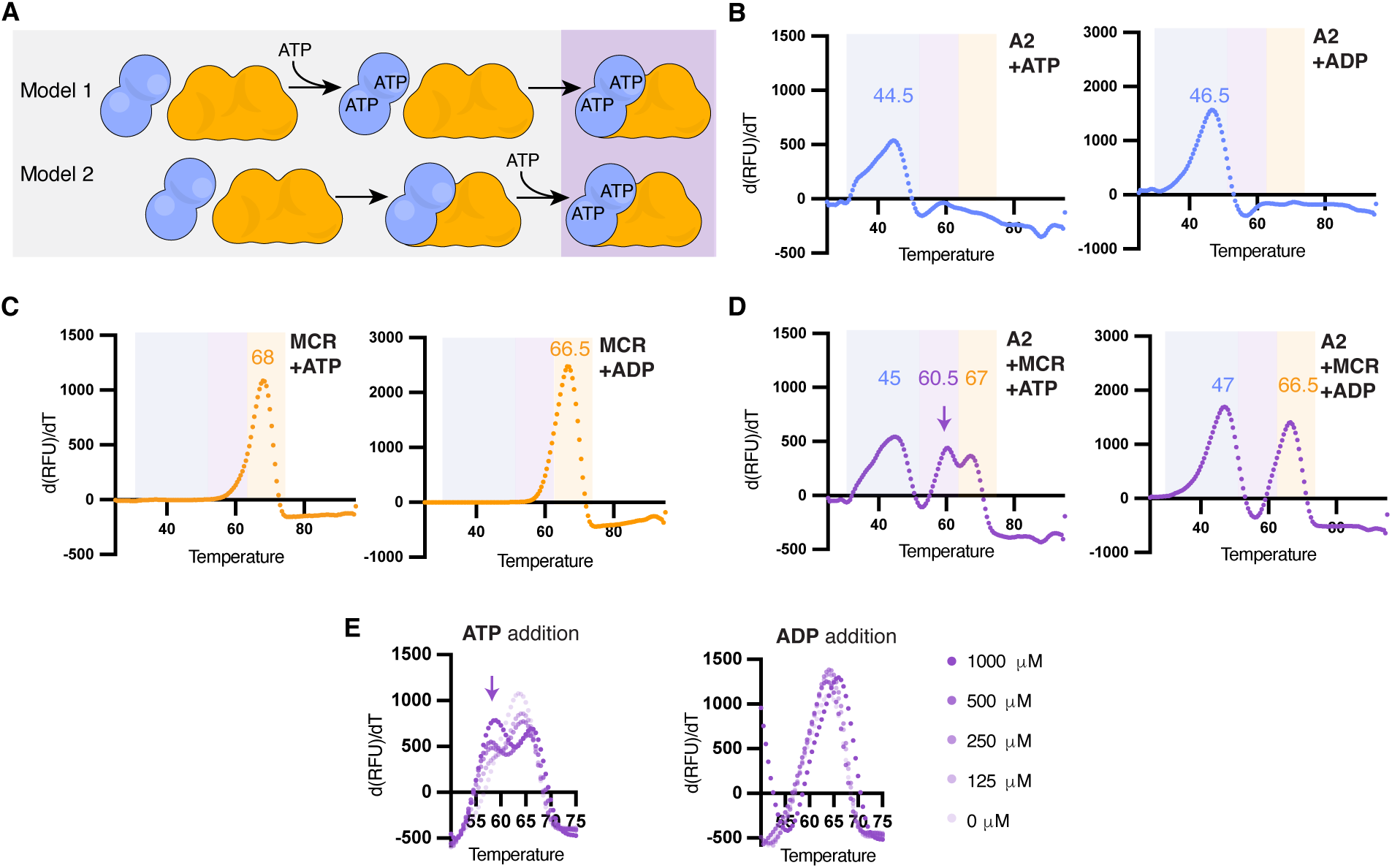
Only ATP-bound component A2 can interact with MCR. (a) Cartoons showing two models of ATP-based interaction between component A2 and MCR. In Model 1, component A2 (blue) binds MCR (in orange) and the component A2-MCR complex interact with ATP. In Model 2, component A2 is bound to ATP (in blue) prior to engagement with MCR (in range). (b-d) Differential scanning fluorimetry (DSF) of (b) component A2, (c) MCR, and (d) component A2 combined with MCR. Samples contained 2 mM ATP (left panel) or ADP (right panel), 10 mM MgCl_2_, 20 mM HEPES, 300 mM NaCl, and 1% glycerol. (e) DSF of component A2 combined with MCR with increasing concentrations as indicated of ATP (left) or ADP (right) and the same assay conditions as (b-d). The peak corresponding to the component A2-MCR complex (purple arrow) only appears in the presence of ATP. All samples in panels b-e contain 0.5 mg/mL of each protein indicated.

AlphaFold (34) predicted structures of apo-component A2 compared to the ATP- and ADP-bound versions reveal significant conformational changes (*SI Appendix*, Fig. S13). Additionally, the AlphaFold predicted structure of ATP-bound component A2 is in high agreement with the MCR-bound component A2 from the activation complex (13) (*SI Appendix*, Fig. S13e). These structural predictions further support our model that only ATP-bound component A2 can interact with MCR (see Model 1 in Fig. 4a).

### A catalytically dead component A2 can still interact with MCR

Based on our model so far, mutations in the Walker A motif of the NBDs that disrupt ATP-binding in component A2 would abolish interaction with MCR. However, whether the Walker B motif of the NBD or the ZBM contributes to the interaction between component A2 and MCR is not as intuitive. To test the role of each of these motifs in MCR interaction, we assessed the amount of MCR that copurifies with each component A2 mutant by immunoblotting against a previously described anti-McrA antibody (26). As expected, the K43A/K329A double mutant, which cannot bind ATP, copurifies with practically no MCR compared to WT (Fig. 5a and *SI Appendix*, Fig. S14). However, each of the single mutants (K43A and K329A) could still interact with MCR but to slightly lesser degree than WT. The E459A, E200A/E459A and C70A/C73A/C86A/C89A mutants co-purified with similar amounts of MCR as WT (Fig. 5a and *SI Appendix*, Fig. S14). However, the E200A mutant had diminished interaction with MCR, like the K43A and K329A mutants. To further corroborate these findings, we performed DSF assays of each of these mutants alone and with MCR in the presence of ATP. No evidence of the component A2-MCR complex could be detected when the K43A/K329A mutant was combined with MCR in the presence of ATP (Fig. 5b and *SI Appendix*, Fig. S15a). In contrast, a peak corresponding to the Tm of the component A2-MCR complex was observed for all the other mutants (Figs. 5c-5h and *SI Appendix*, Figs. S15, S16). These data indicate that the interaction between MCR and component A2 is dependent on ATP-binding but not linked to ATP hydrolysis.

**Figure 5.**
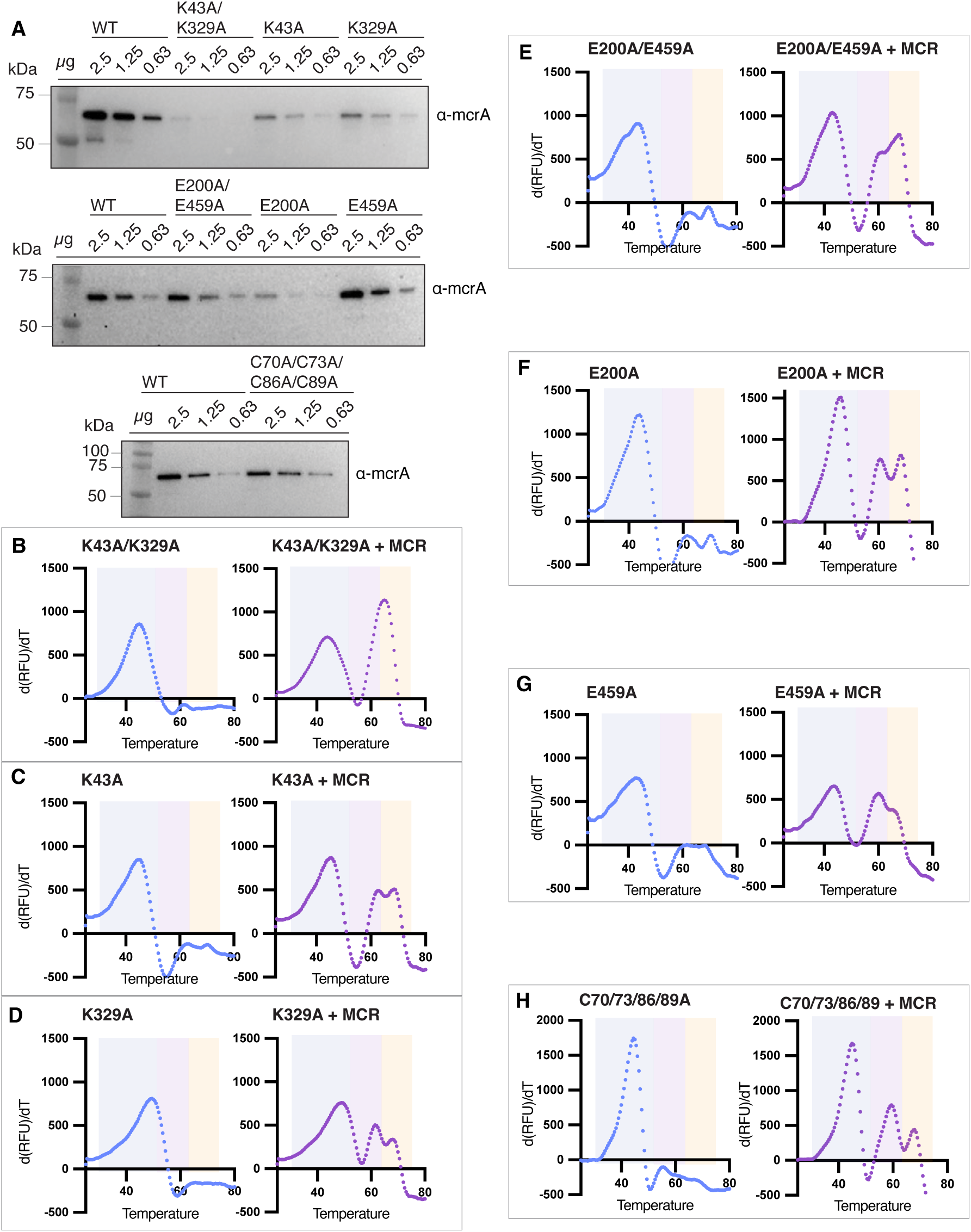
**Interaction of MCR and component A2 is not coupled to ATP hydrolysis.** a) Immunoblots with anti-McrA specific antibodies to evaluate the amount of MCR that copurifies with variants of component A2: K43A/K329A, K43A, and K329A (top), E200A/E459A, E200A, and E459A (middle) and C70A/C73A/C86A/C89A (bottom). Each blot shows a 2X dilution series for each indicated protein starting at 2.5 µg. (b-h) Differential scanning fluorimetry (DSF) of component A2 variants alone (left) or combined with MCR (right). Peaks with a blue background correspond to component A2, with an orange background correspond to MCR, and with a purple background correspond to component A2-MCR complex. Samples contained 2 mM ATP, 10 mM MgCl_2_, 20 mM HEPES, 300 mM NaCl, 0.5 mg/mL of each indicated protein, and 1% glycerol.

### Component A2 has co-evolved with members of the alkyl-coenzyme M reductase (ACR) superfamily

To study the evolutionary history of component A2, we built a phylogenetic tree using 80 sequences that capture the sequence diversity of the protein as well as the taxonomic breadth of archaea that encode them (Fig. 6, Dataset 1). Many archaea, including *M. acetivorans,* also encode a component A2 homolog that contains both NBDs but lacks the ZBM, and we designated these proteins as the outgroup to root the component A2 tree. The phylogeny of component A2 is not consistent with vertical inheritance and indicates that it has undergone rampant HGT within archaea. For example, component A2 sequences from archaea within the Halobacteriota can be found in at least 8 distinct clades that have a bootstrap support of 100 (Fig. 6). *Methanoliparia* encode both MCR and ACR (35) and two distantly related component A2 sequences. Each component A2 sequence from *Methanoliparia* either clusters with sequences derived from archaea that encode MCR or ACR. Methanogens that encode multiple MCR isozymes (36) (members of the Methanobacteriota) only encode one component A2 (Dataset 1) and there are no obvious distinctions between the component A2 sequences derived from ANME and methanogens either. Altogether, these observations suggest that the MCR-specific component A2 is likely to be compatible with all MCRs but not with ACRs and vice versa. Finally, even though component A2 sequences from ACR-encoding archaea are more closely related to each other than to their counterparts from MCR-encoding archaea (Fig. 6), they do not form a highly divergent monophyletic clade like ACRs do relative to MCR (7). This is likely because all component A2 sequences perform the same core function of ATP hydrolysis regardless of which ACR they associate with, in contrast to the significant sequence divergence ACR has undergone to accommodate distinct substrates.

**Figure 6.**
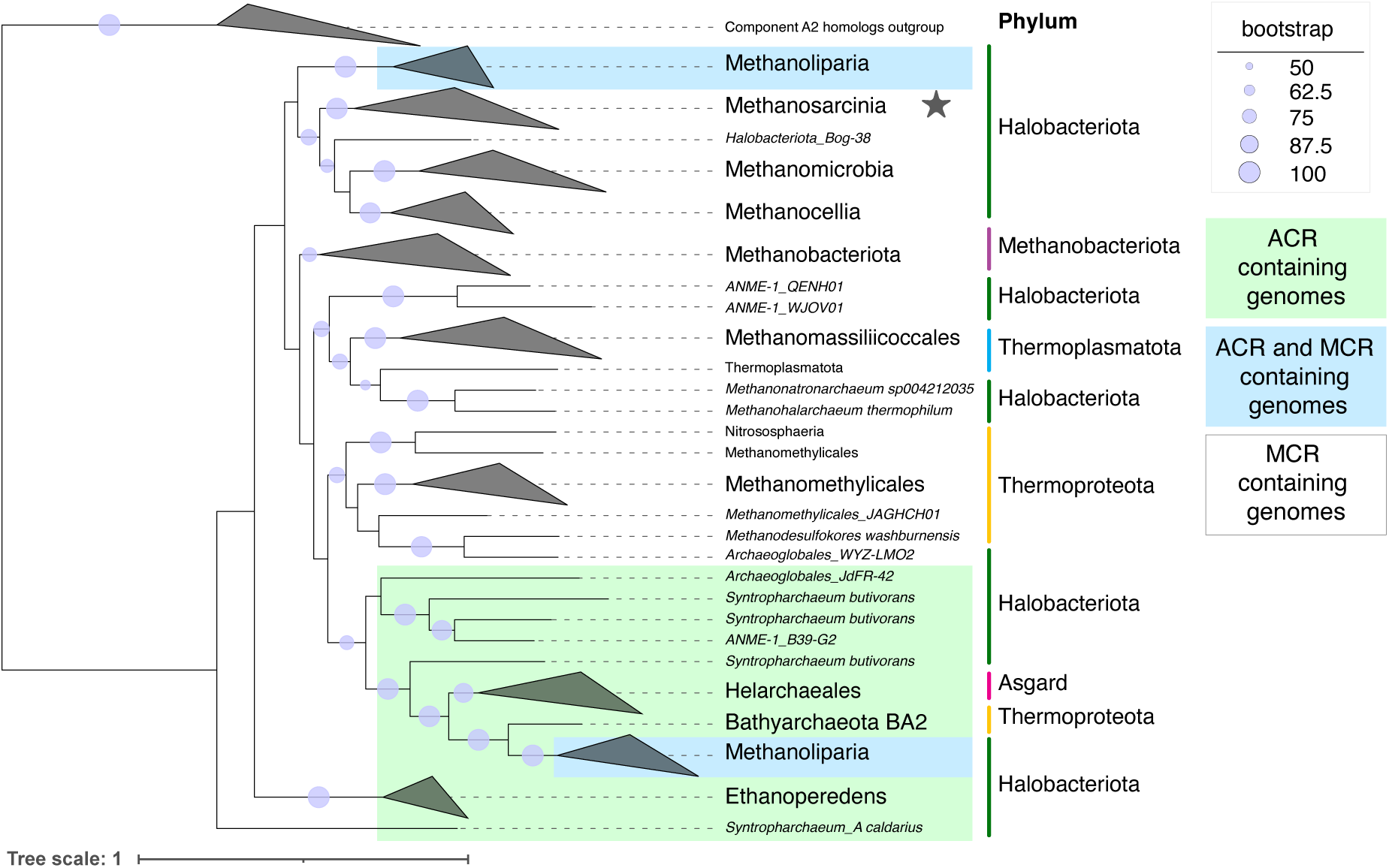
Phylogenetic analysis of component A2 across alkyl-coenzyme M reductase (ACR)-encoding archaea. Phylogenetic tree of component A2 amino acid sequences derived from genomes encoding alkyl coenzyme-M reductase (ACR) in green, methyl-coenzyme M reductase (MCR) in white, and both in blue. All 888 protein sequences with the TIGRFAM Hidden Markov Model (HMM) TIGR0326 assigned to component A2 were obtained from the Genome Taxonomy Database (GTDB) R214. These sequences were clustered at 70% sequence similarity and any truncated sequences or sequences with large insertions (as described in the Methods) were removed, which resulted in a final set of 80 component A2 sequences. The outgroup comprises of 20 sequences of a component A2 homolog lacking a zinc-binding motif (ZBM) that was identified using the sequence from *M. acetivorans* (MA_3967 or MA_RS20695) as a search query. The clade containing the component A2 sequence from *M. acetivorans* is indicated by a star.

## Discussion

In this study, we established an over-expression system in *M. acetivorans* to experimentally evaluate the role of component A2 in the reductive activation of MCR. Many hypotheses regarding the role of component A2 have been made with protein purified from *E. coli* or from indirect measurements using methane production rates as a proxy for ATPase activity. We suspect that the use of heterologously produced protein and an insensitive assay technique have led to confounding results for decades. By directly measuring Pi production by component A2 derived from *M. acetivorans*, we show that this enzyme is a bona fide ATPase, whose activity is dependent on redox conditions (Figs. 2h and 2i) and is modulated by its association with MCR (Fig. 2h). Even though both NBDs of component A2 are required for interaction with MCR and for optimal ATP hydrolysis they are not functionally equivalent (Figs. 3e-j). Mutations in the Walker B motif of NBD2 impede ATP hydrolysis far more than mutations in the Walker B motif of NBD1 (Fig. 3i). In contrast, mutations in the Walker B motif of NBD1 substantially lower component A2-MCR interactions whereas mutations in the Walker B motif of NBD2 do not affect protein-protein interaction at all (Fig. 5a). We hypothesize that the universally conserved ZBM renders component A2 redox-sensitive (Fig. 3c). It is quite likely that the ZBM acts as a “redox-switch” where reduced cysteine residues bind Zn^2+^ and oxidation of these residues reduces rates of ATP hydrolysis (Fig. 3g) (37). This redox switch in component A2 might ensure that the reductive activation of MCR only occurs under environmental conditions that are conducive for catalysis, i.e., in the absence of oxygen or other oxidants.

The evolutionary ties between the maturation machinery for nitrogenases and MCR have been discussed previously (13) but are worth revisiting in consideration of our findings. While nitrogenases requires ATP hydrolysis for electron transfer during catalytic turnover (38), MCR requires ATP input only for cofactor reduction during enzyme maturation. Thus, the net demand for ATP hydrolysis is much lower for MCR than nitrogenases. Furthermore, in nitrogenases, ATP hydrolysis is directly coupled to a redox gating mechanism that triggers electron flow from a [4Fe-4S] cluster in NifH to the P-cluster in NifDK, which is the terminal electron donor for N_2_ reduction (39). Our data suggest that component A2, by itself, is unlikely to be involved in a redox gating mechanism for MCR activation but might facilitate this process through a series of cascading events that are somehow linked to ATP hydrolysis. Our DSF assays reveal a distinct peak for the component A2-MCR complex, which indicates that the two proteins interact even in the absence of McrC and the other MMPs that are putatively involved in transferring electrons to F430 in the active site of MCR (Figs. 4d and 4e). That said, even when an equal amount of MCR is added to component A2, the rate of ATP hydrolysis only increases by 2.2-fold, and the reaction does not proceed to completion (Figs. 2h and 3j). These results suggest that ATP hydrolysis is only triggered when component A2 interacts with a certain sub-population of MCR, likely MCR bound to the electron transfer components of the activation complex. By decoupling MCR interaction from ATP hydrolysis, component A2 might be able to rapidly associate with MCR but only hydrolyze ATP when MCR is primed for reductive activation. Our proposal is also consistent with the mode of action for type I/II ABC transporters where conformational changes in the membrane-bound transporter that are triggered by the interaction of the substrate with the substrate-binding protein ultimately led to hydrolysis by the ATPase component (40). The conformational cues in MCR that are induced by other components of the activation complex to ultimately trigger ATP hydrolysis by component A2 are currently unknown but an exciting area of investigation for future studies.

In summary, this work provides clear experimental evidence that component A2 is an unusual ATPase involved in the activation of MCR. We also establish a pipeline for holistic and mechanistic analyses of essential components involved in the biogenesis of MCR in vivo. A detailed understanding of each component of the activation complex will be necessary for a complete understanding of the intricate pathway involved in the maturation of MCR within the cell.

## Acknowledgements

We thank all members of the Nayak lab for their valuable feedback and support. DDN acknowledges funding from the Searle Scholars Program sponsored by the Kinship Foundation, the Rose Hills Innovator Grant, the Beckman Young Investigator Award sponsored by the Arnold and Mabel Beckman Foundation, the Alfred P. Sloan Research Fellowship sponsored by the Sloan Foundation, the Simons Foundation Early Career Investigator in Marine Microbial Ecology and Evolution Award, the Packard Fellowship in Science and Engineering sponsored by the David and Lucille Packard Foundation, and the Department of Energy through project numbers S589706 and DE-SC0026118. DDN is a Chan-Zuckerberg Biohub – San Francisco Investigator. SAA was supported in part by the NIH-Funded Genetic Dissection of Cells and Organisms Training Program (1T32GM132022-01). The funders had no role in the conceptualization and writing of this manuscript or the decision to submit the work for publication.

## Author contributions

SA was involved in conceptualization, data curation, formal analysis, methodology, investigation, validation, visualization, writing – original draft preparation, and writing – review and editing.

DDN was involved in conceptualization, data curation, funding acquisition, investigation, methodology, project administration, resources, supervision, validation, writing – original draft preparation, and writing – review and editing.

## Data availability statement

Raw Illumina reads associated with genome-wide DNA sequencing have been deposited to the Sequencing Reads Archive (SRA) and will be available upon publication. All other data generated in this study are provided in the manuscript.

## Declaration of interests

The authors declare no competing interests.

## Methods

### Media and culture conditions

All *M. acetivorans* strains were grown at 37 °C without shaking in hermetically sealed Balch tubes or anaerobic bottles in bicarbonate-buffered high-salt (HS) medium containing either 50 mM (for standard passaging) or 100 mM (for protein purification) Trimethylamine.HCl (TMA) as the growth substrate. All *E. coli* strains were grown shaking at 37 °C in lysogeny broth (LB) with 20 µg/mL chloramphenicol.

### Plasmid construction

The vector to overexpress component A2 (MA_3998 or MA_RS20860) in *M.* acetivorans was constructed using the pJK027A vector backbone. Briefly, MA_3998 was amplified from the *M. acetivorans* genome and the 2X Strep- 1X FLAG TAP tag was added to the N-terminus of the protein via primer overhangs. The amplified PCR product was assembled into pJK027A linearized with *NdeI* and *HindIII* via Gibson Assembly as previously described (41). Site directed mutagenesis of component A2 to generate the K43A, K329A, K43A/K329A, E200A, E459A, and E200A/E459A variants was conducted with primers containing the desired point mutations. The C70A/C73A/C86A/C89A allele of A2 was generated as a synthetic construct (Twist Biosciences, South San Francisco, CA, USA). All plasmids were transformed into *E. coli* strain WWM4489 (42), a derivative of DH10ß, by electroporation (MicroPulser Electroporator, Bio-Rad, Hercules, CA, USA) and the resulting transformants were grown in LB supplemented with 20 µg/mL chloramphenicol and10 mM rhamnose to induce high copy number of the plasmid (42). All plasmids were extracted using the Zyppy Miniprep Kit (Zymo Research, Tustin, CA) and verified by Sanger sequencing at the UC Berkeley DNA Sequencing Facility. All primers used were obtained from Integrated DNA Technologies, Coralville, IA, USA. All primers and plasmids used in this study can be found in *SI Appendix*, Tables S4 and S5.

### Transformation of Methanosarcina acetivorans

*M. acetivorans* strain WWM73 (29) was transformed with the component A2 overexpression plasmids by liposome-mediated transformation as previously described (43). Each transformation reaction was performed with 2 µg plasmid DNA and 20 mL of *M. acetivorans* cells grown to late-exponential phase high-salt (HS) minimal medium with 50 mM trimethylamine.hydrochloride (TMA) as the sole carbon and energy source containing HS-media. Transformants were then plated on agar-solidified HS-medium with 50 mM TMA and 2 µg/mL puromycin. Plates were incubated at 37 °C for 2-3 weeks in an intrachamber anaerobic incubator with an H_2_S /CO_2_/N_2_ (1,000 ppm/20%/balance) headspace. Colonies were picked into 10 mL of HS-medium with 50 mM TMA and 2 µg/mL puromycin and incubated at 37 °C without shaking. Once grown, integration of the plasmid onto the chromosome was confirmed by PCR using diagnostic primers listed *SI Appendix*, Table S4. PCR product sequences were verified by Sanger sequencing at the UC Berkeley DNA Sequencing Facility. Strains generated for this study are listed in *SI Appendix*, Table S6.

### Genomic DNA extraction and whole-genome sequencing

Genomic DNA was extracted from 2 mL of culture grown to saturation using the QIAGEN DNeasy Blood & Tissue kit (QIAGEN, Hilden, Germany) according to manufacturer’s protocol. Library preparation and Illumina sequencing was conducted by SeqCenter (Pittsburgh, PA, USA). The resulting sequencing reads were analyzed *breseq* v0.35.5 (44) using default settings.

### Growth curves of Methanosarcina acetivorans

Growth curves were performed by preculturing strains in HS-TMA media without shaking at 37 °C (HeraTherm General Protocol Microbiological Incubator, Thermo Fisher Scientific, Waltham, MA) in sealed Balch tubes. From early stationary phase cells, 0.5 mL culture was transferred into three replicate tubes with 100 µg/mL tetracycline and three replicate tubes without tetracycline. All tubes contained 2 µg/mL puromycin to maintain the integrated plasmid. Growth was measured by monitoring the optical density at 600 nm (Genesys 50, Thermo Fisher Scientific, Waltham, MA). All tubes containing tetracycline were wrapped in aluminum foil to prevent light-based degradation of the chemical. The growth rate was calculated from the slope of the linear fit of the log_10_-transformed optical density versus time with a minimum five points in the exponential phase with the highest R^2^ value (minimum cut-off ≥ 0.99).

### Anaerobic purification of proteins

500 mL cultures of *M. acetivorans* were grown in HS-media supplemented with 100 mM TMA in 1-liter anaerobic bottles sealed with butyl rubber stoppers (Chemglass Life Sciences, Vineland, NJ, USA) at 37 °C. Cultures were supplemented with 2 µg/mL puromycin and 100 µg/mL tetracycline to induce protein expression and grown to late-exponential phase (for component A2) or stationary phase (for methyl-coenzyme M reductase; MCR). All the steps of protein purification were conducted under anaerobic conditions in a Coy chamber (Coy Lab Products, Grass Lake, MI, USA) with a N_2_/H_2_ (96-97%/ 3-4%) headspace. All buffers were made anaerobic by sparging with 100% N_2_ for 30 minutes and were filter sterilized following sparging. Cells were harvested by spinning (Sorvall Legend XTR, Thermo Fisher Scientific, Waltham, MA, USA) at 6000 RPM at 4 °C for 10 minutes in a gas-tight sealed polypropylene Nalgene centrifuge bottles. After harvesting, cells were anaerobically lysed via osmotic pressure in 20 mM HEPES, 1% glycerol buffer, pH 8 (Buffer A) with the addition of 0.25X Roche cOmplete EDTA-free Protease Inhibitor Cocktail (MilliporeSigma, Burlington, MA, USA) and DNase I (Thermo Fisher Scientific, Waltham, MA, USA). After lysis, 5 M NaCl was added to crude cell lysate for a final concentration of 300 mM NaCl. Crude cell lysate was cleared by spinning at 10,000 RPM at 4 °C for 30 minutes. To purify protein, cleared cell lysate was applied twice to 0.25-0.5 mL Strep-Tactin Superflow plus resin (Qiagen, Hilden, Germany) equilibrated with 10 mL anaerobic Buffer A + 300 mM NaCl. After washing the resin with 12 mL of Buffer A + 300 mM NaCl, the protein of interest was eluted with Buffer A + 300 mM NaCl + 2.5 mM desthiobiotin (Sigma-Aldrich, St. Louis, MO, USA). Purified protein was stored at room temperature in the anaerobic chamber, unless otherwise stated (45). Protein concentration was determined by Pierce Bradford Plus Protein Assay Reagent (Thermo Fisher Scientific, Waltham, MA, USA) using Bovine Serum Albumin (BSA) for calibration curve generation.

### ATPase assays

All ATPase assays were performed under anaerobic conditions in the Coy anaerobic chamber unless otherwise stated. Assays were performed with component A2 within 48 hours of purification. Component A2 was stored anaerobically at room temperature (RT) prior to use (ref). ATPase reactions were performed in anaerobically in 110 µL reaction volume incubated at 37 °C containing 20 mM HEPES, 1% glycerol, 300 mM NaCl, 10 mM MgCl_2_, 200 µM ATP, and 500 µg/mL of either component A2, MCR, or both. Reactions were started with addition of protein. Reactions were set up in technical triplicate and 25 µL aliquots were taken at 0, 15, 30, and 60-minute time points. To quench reaction time points, aliquots were immediately added to an equivalent volume of ice cold 20 mM EDTA, pH 8, and removed from the anaerobic chamber. Aliquots were stored on ice until the final time point was taken. ATPase activity was evaluated immediately following the final time point by measuring the production of inorganic phosphate using malachite green solution (Echelon Biosciences, Salt Lake City, UT, USA). Each 50 µL aliquot was plated in duplicate in a 96-well plate and 80 µL of malachite green solution was added to each well and mixed by pipetting. After allowing color development for 25 minutes at RT, the absorbance was measured at 620 nm on an Epoch 2 microplate reader (BioTek, Winooski, VT, USA). Absorbance was converted to phosphate concentration using a 0-100 µM phosphate calibration curve and plotted against time to determine ATPase activity over the course of the reaction. Reactions were internally normalized to the 0-minute time point to only measure change in phosphate production over the course of the 60-minute incubation.

### Differential Scanning Fluorimetry

Anaerobically purified component A2 or MCR was removed from the anaerobic chamber to set up reactions for differential scanning fluorimetry at RT on the bench top. Each sample contained 500 µg/mL indicated protein, 20 mM MgCl_2,_ ATP at concentration indicated in figure legend (ranging from 0 – 2 mM), 20 mM HEPES, 300 mM NaCl, and 1% glycerol at pH 8. SYPRO Orange Protein Gel Stain (Invitrogen, Thermo Fisher Scientific, Waltham, MA, USA) was added to a final concentration of 5X and each sample was gently pipetted to mix and dispensed into 96-well PCR plates (Applied Biosystems, Thermo Fisher Scientific, Waltham, MA, USA) and sealed with adhesive optical film. Using a CFX96 Touch Real-Time PCR Detection System (Bio-Rad, Hercules, CA, USA) fluorescence was measured in the FRET channel (450-730 nm range of excitation/emission wavelengths) along a temperature gradient from 25°C to 95°C in increments of 0.5°C every 30 seconds. The first derivative of the fluorescence emission as a function of temperature (dRFU/dT) was plotted and the melting temperature (T_m_) was defined as the peak(s) of the first derivative. Melt curves were repeated with biological triplicate (independent protein purifications) as indicated.

### SDS-PAGE and Immunoblotting of FLAG-tagged proteins and of MCR

Samples containing protein as indicated in figure captions were combined with Laemmli sample buffer (Bio-Rad, Hercules, CA, USA) and 2.5% β-mercaptoethanol and then heated at 95°C for 8 minutes. Samples were then loaded into 12% precast Tris-Glycine denaturing SDS-PAGE gels (Mini-PROTEAN TGX, Bio-Rad, Hercules, CA, USA). Bio-Rad Precision Plus Prestained Protein Standard was used as a molecular weight standard. Gels were run at 100-150 V until the dye front reached the bottom of the cassette. Gels were stained with Coomassie GelCode Blue (Thermo Fisher Scientific, Waltham, MA, USA). For immunoblotting, protein was transferred from the gel to the PVDF (polyvinylidene fluoride) membrane with the Trans-Blot Turbo Transfer System and Midi-PVDF Transfer Pack (Bio-Rad, Hercules, CA, USA) following manufacturer’s instructions. Following transfer, membranes were rinsed for 5 minutes with water, 5 minutes with PBS (137 mM NaCl, 2.7 mM KCl, 10 mM Na_2_HPO_4_, 1.8 mM KH_2_PO_4_), and blocked for 1 hour at RT with 5% nonfat milk dissolved in PBS. For visualization of FLAG-tagged proteins, membranes were then rinsed in PBS 4X for 5 minutes prior to incubation with mouse monoclonal anti-FLAG M2 HRP-conjugated antibody (Sigma-Aldrich, St Louis, MO) diluted 1:66666 in PBS-T (PBS with 0.05% Tween-20) for 1 hour at RT. Following incubation with antibody, membranes were rinsed 3X in PBS-T and then 3X in PBS for 5 minutes each. Alternatively, for visualizing McrA, membranes were rinsed in PBS 4X for 5 minutes following blocking and then incubated overnight at 4°C with polyclonal rabbit antibodies raised against McrA (1:10000 dilution) (GenScript, Piscataway, NJ, USA) in PBS-T. The following day membranes were rinsed in PBS-T 4X for 5 minutes each and then incubated with anti-rabbit horseradish peroxidase (HRP) conjugate antibodies (1:100000 dilution) (Promega, Madison, WI, USA) for 2 hours at room temperature and finally rinsed 3X in PBS-T and then 3X in PBS for 5 minutes each. For both FLAG and McrA visualization, the signal was developed by addition of the Immobilon Western Chemiluminescent HRP Substrate (MilliporeSigma, Burlington, MA, USA) in the dark. Both SDS-PAGE gels and immunoblots were visualized on a ChemiDoc MP Imaging System (Bio-Rad, Hercules, CA, USA).

### Mass spectrometry Sample Identification

Mass spectrometry-based identification of proteins extracted from SDS-PAGE gels were performed by either Applied Biomics, Inc (Hayward, CA, USA) or QB3/Chemistry Mass Spectrometry Facility at UC Berkeley. Applied Biomics performed sample preparation in house. For samples analyzed at the UC Berkeley QB3/Chemistry Mass Spectrometry Facility sample preparation was performed as follows. Bands from Coomassie stained SDS-PAGE gel were excised with a razor blade and diced, and gel pieces were washed for 20 minutes in 500 µL of 100 mM NH_4_HCO_3_ and the supernatant was discarded. Gel pieces were then incubated in 150 µL of 100 mM NH_4_HCO_3_ and 10 µL of 45 mM DTT for 15 minutes at 50 °C. Next 10 µL of 100 mM iodoacetamide was added and the mixture was incubated for 15 minutes in the dark at RT. The supernatant was discarded gel pieces were washed with 500 µL of a 50:50 mix of acetonitrile and 100mM NH_4_HCO_3_ with shaking for 20 minutes. After the supernatant was discarded, gel pieces were incubated with 50 µL of acetonitrile for 15 minutes. Solvent was removed and gel fragments were dried in a speed vac. Gel pieces were then reswelled with 10 µL of 25 mM NH_4_HCO_3_ containing 0.1 µg sequencing-grade modified trypsin (Promega, Madison, WI, USA). After 15 minutes, 20 µL of additional NH_4_HCO_3_ was added and the reaction was incubated overnight at 37 °C. Remaining peptides from the gel pieces were extracted twice with 50 µl of 60% acetonitrile/0.1% formic acid for 20 minutes, then once with 25 µl acetonitrile, and samples were dried in a speed-vac.

### RNA extraction and cDNA synthesis

*M. acetivorans* strain WWM60 was grown to mid-exponential phase and 1 mL of culture was added to an equivalent volume of Trizol (Life Technologies, Carslbad, CA) prewarmed to 37 °C. Following incubation for 5 minutes at RT, 2 mL of 100% cold ethanol was added, and RNA was extracted according to the manufacturer’s instructions using the Qiagen RNeasy Mini Kit (Qiagen, Hilden, Germany). Concentration of the RNA was determined using a Nanodrop One UV Spectrophotometer (Thermo Fisher Scientific, Waltham, MA) and stored at −80 °C. To eliminate any minor amounts of contaminating genomic DNA, RNA was treated with DNase according to the manufacturer’s instructions using TURBO-DNA free kit (Thermo Fisher Scientific, Waltham, MA, USA). Next, to synthesize cDNA, reactions were set up with 2.5 ng/µL random hexamers, 10 ng/µL DNase-treated RNA, 0.5 mM dNTPs, 5 mM DTT, First-strand buffer, and 10 U/µL Superscript III. Prior to the addition of DTT, First-strand buffer, and Superscript III, the reaction was incubated at 65 °C for 5 minutes and then placed on ice for 1 minute. Following the addition of the rest of the reagents, the reaction was incubated at 25 °C for 5 minutes, then 50 °C for 60 minutes, and finally 70 °C for 15 minutes. The resulting cDNA was stored at -20°C. All reagents used for cDNA synthesis were obtained from Thermo Fisher Scientific, Waltham, MA, USA.

### Protein structural prediction and visualization

AlphaFold 3 (34) was used for structural models of *M. acetivorans* component A2 (MA_3998) alone and with ATP or ADP. Protein domains were identified using InterPro v108 (46). Protein structures were visualized using ChimeraX v1.7.1 (47).

### Protein sequence alignment

Protein sequences with the assigned TIGRFAM Hidden Markov Model (HMM) (48) TIGR03269 were downloaded from the Genome Taxonomy Database (GTDB) R214, which included 888 sequences. A multiple sequence alignment (MSA) MUSCLE v5.1 alignment (49) was performed using Geneious software v 2024.0.2. The MSA of individual motifs of interest (NBD1 walker-A, NBD2 walker-A, NBD1 walker-B, NBD2 walker-B, and zinc binding motif) were extracted from the full sequence alignment. Each individual motif contained the following number of sequences from the MSA: 867 in NBD1 walker-A, 888 in NBD2 walker-A, 883 in NBD1 walker-B, 880 in NBD2 walker-B, and 870 in the zinc binding motif. Individual sequence motifs were visualized with WebLogo v3 (50).

### Phylogenetic analysis

All 888 sequences were clustered based on 70% sequence similarity using Cluster Database at High Identity with Tolerance (CD-HIT) (51) with default parameters. Clustering provided a list of 86 sequences that were further trimmed based on sequence length to omit truncated sequences or sequences with large insertions. Sequences were retained if they fell within ±10% of the mean sequence length, and the component A2 sequence from *M. acetivorans* was manually added in, due to its relevance to this work. This resulted in a final set of 80 sequences. The tree was built using these 80 sequences along with 20 sequences for the outgroup. The outgroup was determined by using Basic Local Alignment Search Tool (BLAST) to find 20 sequences that are most like the component A2 homolog from *M. acetivorans* MA_3967 (MA_RS20695). A multiple sequence alignment (MSA) was performed using the MUSCLE v5.1 plug-in (49) in Geneious software v2024.0.2. A tree was then built using the Randomized Axelerated Maximum Likelihood (RAxML) algorithm v8.2.11 (52) with protein model GAMMABLOSUM62 and 100 bootstrap replicates. The tree was visualized using iTOL v7 (Interactive Tree of Life) (53).

## Notes

### Competing Interest Statement

The authors have declared no competing interest.

## References

1. R. K. Thauer, A.-K. Kaster, H. Seedorf, W. Buckel, R. Hedderich, Methanogenic archaea: ecologically relevant differences in energy conservation. Nat Rev Microbiol 6, 579–591 (2008).

2. A. Gendron, K. D. Allen, Overview of Diverse Methyl/Alkyl-Coenzyme M Reductases and Considerations for Their Potential Heterologous Expression. Frontiers in Microbiology 13, 867342 (2022).

3. S. J. Hallam, et al., Reverse Methanogenesis: Testing the Hypothesis with Environmental Genomics. Science 305, 1457–1462 (2004).

4. B. Ginovska, S. Raugei, S. W. Ragsdale, C. Ohmer, R. Sarangi, Structural and Mechanistic Advances in the Chemistry of Methyl-Coenzyme M Reductase (MCR). Acc. Chem. Res. 58, 824–833 (2025).

5. U. Ermler, W. Grabarse, S. Shima, M. Goubeaud, R. K. Thauer, Crystal Structure of Methyl-Coenzyme M Reductase: The Key Enzyme of Biological Methane Formation. Science 278, 1457–1462 (1997).

6. S. Shima, et al., Structure of a methyl-coenzyme M reductase from Black Sea mats that oxidize methane anaerobically. Nature 481, 98–101 (2011).

7. O. N. Lemaire, T. Wagner, A Structural View of Alkyl-Coenzyme M Reductases, the First Step of Alkane Anaerobic Oxidation Catalyzed by Archaea. Biochemistry 61, 805–821 (2022).

8. R. Laso-Pérez, et al., Anaerobic Degradation of Non-Methane Alkanes by “ *Candidatus* Methanoliparia” in Hydrocarbon Seeps of the Gulf of Mexico. mBio 10, e01814–19 (2019).

9. S. Rospert, R. Böcher, S. P. J. Albracht, R. K. Thauer, Methyl-coenzyme M reductase preparations with high specific activity from H _2_ -preincubated cells of *Methanobacterium thermoautotrophicum*. FEBS Letters 291, 371–375 (1991).

10. P. E. Rouvière, T. A. Bobik, R. S. Wolfe, Reductive activation of the methyl coenzyme M methylreductase system of Methanobacterium thermoautotrophicum delta H. J Bacteriol 170, 3946–3952 (1988).

11. W. L. Ellefson, W. B. Whitman, R. S. Wolfe, Nickel-containing factor F430: chromophore of the methylreductase of Methanobacterium. Proc. Natl. Acad. Sci. U.S.A. 79, 3707–3710 (1982).

12. C. Holliger, A. J. Pierik, E. J. Reijerse, W. R. Hagen, A spectroelectrochemical study of factor F430 nickel(II/I) from methanogenic bacteria in aqueous solution. J. Am. Chem. Soc. 115, 5651–5656 (1993).

13. F. Ramírez-Amador, et al., Structure of the ATP-driven methyl-coenzyme M reductase activation complex. Nature (2025). 10.1038/s41586-025-08890-7.

14. S. A. Adler, G. L. Chadwick, D. D. Nayak, Assembly and maturation of methyl-coenzyme M reductase in methanogenic archaea. Current Opinion in Microbiology 87, 102637 (2025).

15. R. K. Thauer, Methyl (Alkyl)-Coenzyme M Reductases: Nickel F-430-Containing Enzymes Involved in Anaerobic Methane Formation and in Anaerobic Oxidation of Methane or of Short Chain Alkanes. Biochemistry 58, 5198–5220 (2019).

16. R. P. Gunsalus, R. S. Wolfe, Methyl coenzyme M reductase from Methanobacterium thermoautotrophicum. Resolution and properties of the components. Journal of Biological Chemistry 255, 1891–1895 (1980).

17. P. E. Rouvière, J. C. Escalante-Semerena, R. S. Wolfe, Component A2 of the methylcoenzyme M methylreductase system from Methanobacterium thermoautotrophicum. J Bacteriol 162, 61–66 (1985).

18. D. P. Nagle, R. S. Wolfe, Component A of the methyl coenzyme M methylreductase system of Methanobacterium: resolution into four components. Proc. Natl. Acad. Sci. U.S.A. 80, 2151–2155 (1983).

19. D. Prakash, Y. Wu, S.-J. Suh, E. C. Duin, Elucidating the Process of Activation of Methyl-Coenzyme M Reductase. J Bacteriol 196, 2491–2498 (2014).

20. K. Zheng, P. D. Ngo, V. L. Owens, X.-P. Yang, S. O. Mansoorabadi, The biosynthetic pathway of coenzyme F430 in methanogenic and methanotrophic archaea. Science 354, 339–342 (2016).

21. S. J. Moore, et al., Elucidation of the biosynthesis of the methane catalyst coenzyme F430. Nature 543, 78–82 (2017).

22. E. B. Shelton, et al., An Expanded Molecular Model for the Activation of Methyl-Coenzyme M Reductase. Biochemistry 64, 4424–4436 (2025).

23. G. Borrel, et al., Wide diversity of methane and short-chain alkane metabolisms in uncultured archaea. Nat Microbiol 4, 603–613 (2019).

24. K. E. Shalvarjian, et al., Methanogenic archaea encoding Pyrrolysine maintain ambiguous amber codon usage. Proceedings of the National Academy of Sciences 122, e2517473122 (2025).

25. B. E. Downing, D. Gupta, K. E. Shalvarjian, D. D. Nayak, Genus-specific remodeling of carbon and energy metabolism facilitates acetoclastic methanogenesis in Methanosarcina spp. and Methanothrix spp. Journal of Bacteriology 208, e00448–25 (2026).

26. G. L. Chadwick, G. A. Dury, D. D. Nayak, Physiological and transcriptomic response to methyl-coenzyme M reductase limitation in Methanosarcina acetivorans. Applied and Environmental Microbiology 90, e0222023 (2024).

27. D. D. Nayak, W. W. Metcalf, Cas9-mediated genome editing in the methanogenic archaeon Methanosarcina acetivorans. Proceedings of the National Academy of Sciences of the United States of America 114, 2976–2981 (2017).

28. J. Saini, A. Dhamad, A. Muniyasamy, A. J. Alverson, D. J. Lessner, The nitrogenase cofactor biogenesis enzyme NifB is essential for the viability of methanogens. [Preprint] (2023). Available at: http://biorxiv.org/lookup/doi/10.1101/2023.10.20.563283 [Accessed 11 September 2024].

29. A. M. Guss, M. Rother, J. K. Zhang, G. Kulkkarni, W. W. Metcalf, New methods for tightly regulated gene expression and highly efficient chromosomal integration of cloned genes for *Methanosarcina* species. Archaea 2, 193–203 (2008).

30. C. H. Kuhner, B. D. Lindenbach, R. S. Wolfe, Component A2 of methylcoenzyme M reductase system from Methanobacterium thermoautotrophicum delta H: nucleotide sequence and functional expression by Escherichia coli. J Bacteriol 175, 3195–3203 (1993).

31. P. I. Hanson, S. W. Whiteheart, AAA+ proteins: have engine, will work. Nat Rev Mol Cell Biol 6, 519–529 (2005).

32. C. Orelle, O. Dalmas, P. Gros, A. Di Pietro, J.-M. Jault, The Conserved Glutamate Residue Adjacent to the Walker-B Motif Is the Catalytic Base for ATP Hydrolysis in the ATP-binding Cassette Transporter BmrA. Journal of Biological Chemistry 278, 47002–47008 (2003).

33. M. F. Khan, et al., Determination of Protein–Ligand Binding Affinities by Thermal Shift Assay. ACS Pharmacol. Transl. Sci. 7, 3096–3107 (2024).

34. J. Abramson, et al., Accurate structure prediction of biomolecular interactions with AlphaFold 3. Nature 630, 493–500 (2024).

35. Z. Zhou, et al., Non-syntrophic methanogenic hydrocarbon degradation by an archaeal species. Nature 601, 257–262 (2022).

36. L. G. Bonacker, S. Baudner, E. Mörschel, R. Böcher, R. K. Thauer, Properties of the two isoenzymes of methyl-coenzyme M reductase in Methanobacterium thermoautotrophicum. Eur J Biochem 217, 587–595 (1993).

37. N. J. Pace, E. Weerapana, Zinc-Binding Cysteines: Diverse Functions and Structural Motifs. Biomolecules 4, 419 (2014).

38. J. B. Howard, D. C. Rees, Structural Basis of Biological Nitrogen Fixation. Chem. Rev. 96, 2965–2982 (1996).

39. H. L. Rutledge, F. A. Tezcan, Electron Transfer in Nitrogenase. Chem Rev 120, 5158–5193 (2020).

40. A. J. Rice, A. Park, H. W. Pinkett, Diversity in ABC transporters: Type I, II and III importers. Crit Rev Biochem Mol Biol 49, 426–437 (2014).

41. D. G. Gibson, et al., Enzymatic assembly of DNA molecules up to several hundred kilobases. Nat Methods 6, 343–345 (2009).

42. S. Y. Kim, et al., Different biosynthetic pathways to fosfomycin in Pseudomonas syringae and Streptomyces species. Antimicrob Agents Chemother 56, 4175–4183 (2012).

43. W. W. Metcalf, J. K. Zhang, E. Apolinario, K. R. Sowers, R. S. Wolfe, A genetic system for Archaea of the genus Methanosarcina: liposome-mediated transformation and construction of shuttle vectors. Proc Natl Acad Sci U S A 94, 2626–2631 (1997).

44. D. E. Deatherage, J. E. Barrick, Identification of mutations in laboratory evolved microbes from next-generation sequencing data using breseq. Methods Mol Biol 1151, 165–188 (2014).

45. E. C. Duin, D. Prakash, C. Brungess, “Methyl-Coenzyme M Reductase from Methanothermobacter marburgensis” in Methods in Enzymology, (Elsevier, 2011), pp. 159–187.

46. M. Blum, et al., InterPro: the protein sequence classification resource in 2025. Nucleic Acids Research 53, D444–D456 (2025).

47. E. C. Meng, et al., UCSF ChimeraX: Tools for structure building and analysis. Protein Science 32, e4792 (2023).

48. D. H. Haft, TIGRFAMs: a protein family resource for the functional identification of proteins. Nucleic Acids Research 29, 41–43 (2001).

49. R. C. Edgar, Muscle5: High-accuracy alignment ensembles enable unbiased assessments of sequence homology and phylogeny. Nat Commun 13, 6968 (2022).

50. G. E. Crooks, G. Hon, J.-M. Chandonia, S. E. Brenner, WebLogo: A Sequence Logo Generator: Figure 1. Genome Res. 14, 1188–1190 (2004).

51. W. Li, A. Godzik, Cd-hit: a fast program for clustering and comparing large sets of protein or nucleotide sequences. Bioinformatics 22, 1658–1659 (2006).

52. A. Stamatakis, RAxML version 8: a tool for phylogenetic analysis and post-analysis of large phylogenies. Bioinformatics 30, 1312–1313 (2014).

53. I. Letunic, P. Bork, Interactive Tree Of Life (iTOL) v5: an online tool for phylogenetic tree display and annotation. Nucleic Acids Res 49, W293–W296 (2021).

