## Supplemental Files for "Component A2 is a redox-sensitive archaeal ATPase activated by methyl-coenzyme M reductase"

### Supplemental Figures

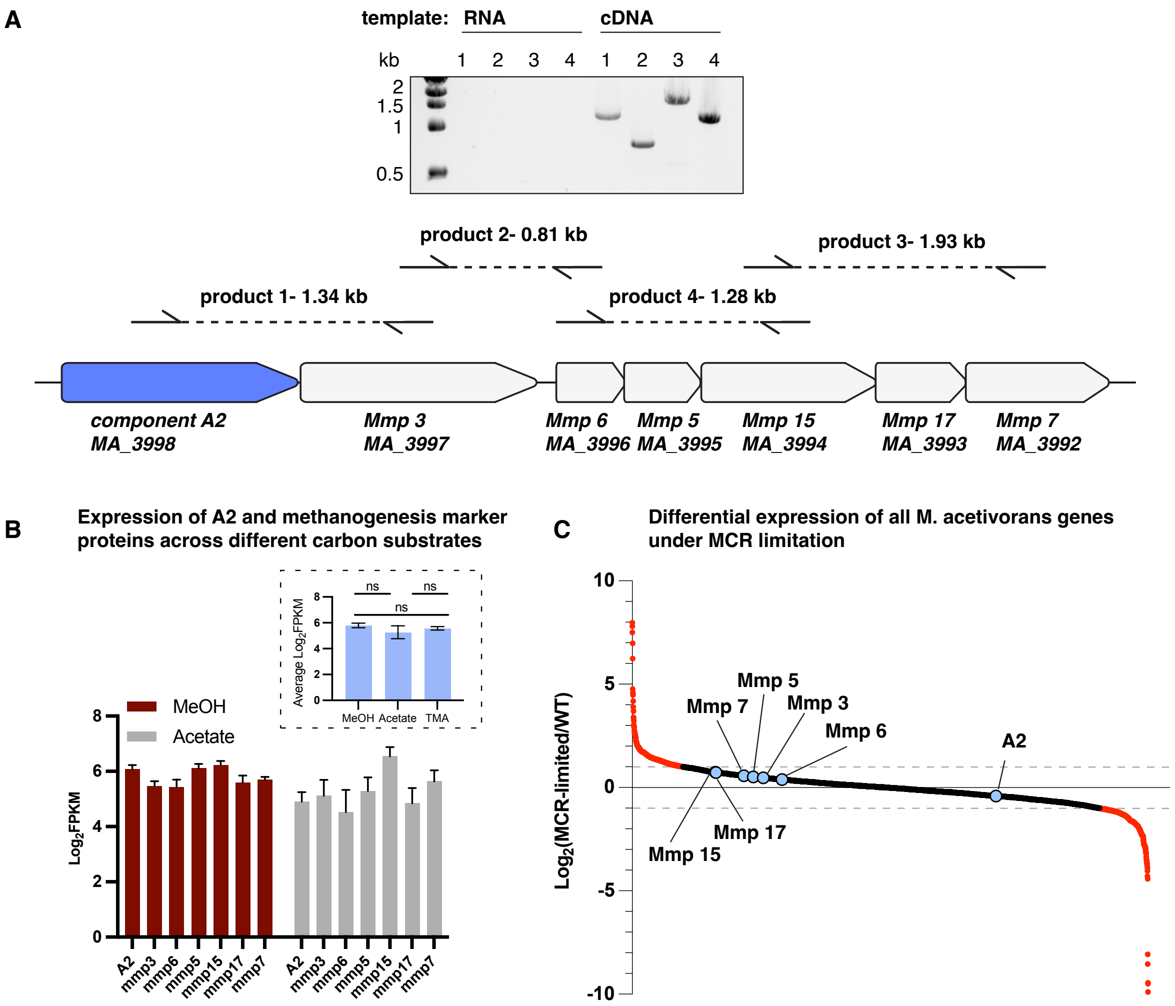


**Fig S1.** Expression analysis of the methyl-coenzyme M reductase (MCR) activation operon in *Methanosarcina acetivorans*. (a) Agarose gel electrophoresis showing the PCR products generated from the primer pairs across all gene boundaries for the MCR activation operon with either RNA or cDNA synthesized from the RNA as the template used for the PCR reactions. Lanes 1-4 correspond to the products for the primers pairs labelled 1-4 on the chromosome map shown below. Component A2 is highlighted in blue. (b) expression of genes in the MCR activation operon on during growth on high-salt (HS) minimal-medium supplemented with 125 mM methanol (MeOH; maroon) or 40 mM acetate (gray) as the sole carbon and energy substrate at 37 °C. Error bars represent the standard deviation for three replicates. Inset shows average expression of the MCR activation operon on MeOH, TMA, and acetate. Error bars represent the standard deviation of three replicates. Statistical significance tested by unpaired Welch's t test. ns indicates a p-value > 0.05 (c) Differential expression of all genes in *M. acetivorans* between strains WWM60 (parent) and DDN032 (P*mcrB*(tetO1)-*mcrBDCGA*) (1) grown without tetracycline to curtail growth by limiting the level of MCR in the cell. Genes are ordered along the x-axis from highest to lowest differential expression. Genes with a log2-fold change > 1 or <1 are show in red and genes with a log2-fold change in expression between -1 and 1 are shown in black. Genes in the MCR activation operon are highlighted.

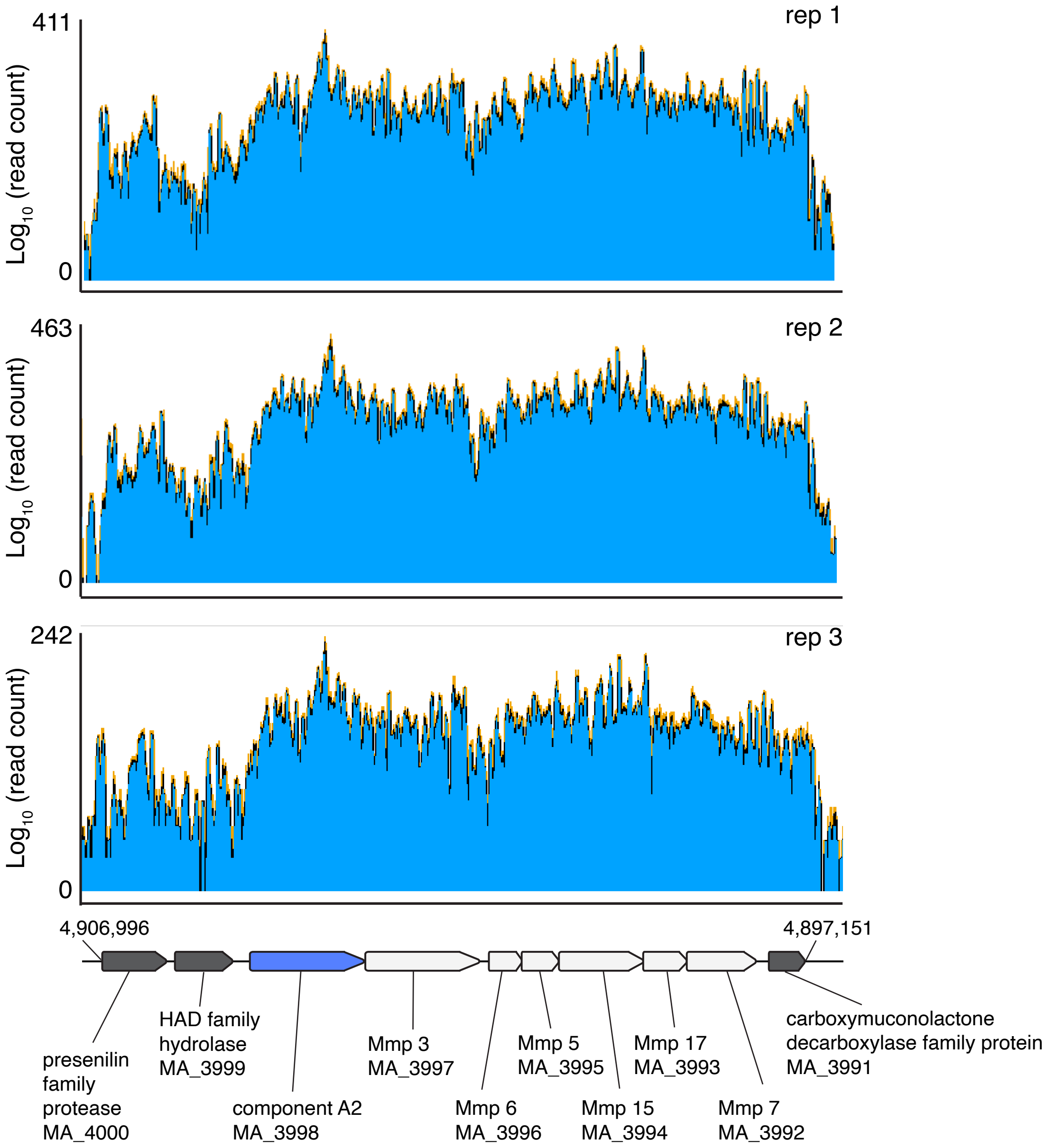


**Fig S2.** RNAseq reads (2) mapped across the genomic region of *M. acetivorans* containing the MCR activation operon. Paired-end RNA read coverage of the MCR activation operon during growth on TMA and ~2 kilobase up stream and ~1 kilobase downstream genes included. Three technical replicates (denoted as rep1, rep2, and rep3) are shown. Reads are aligned to the chromosomal locus as indicated below.

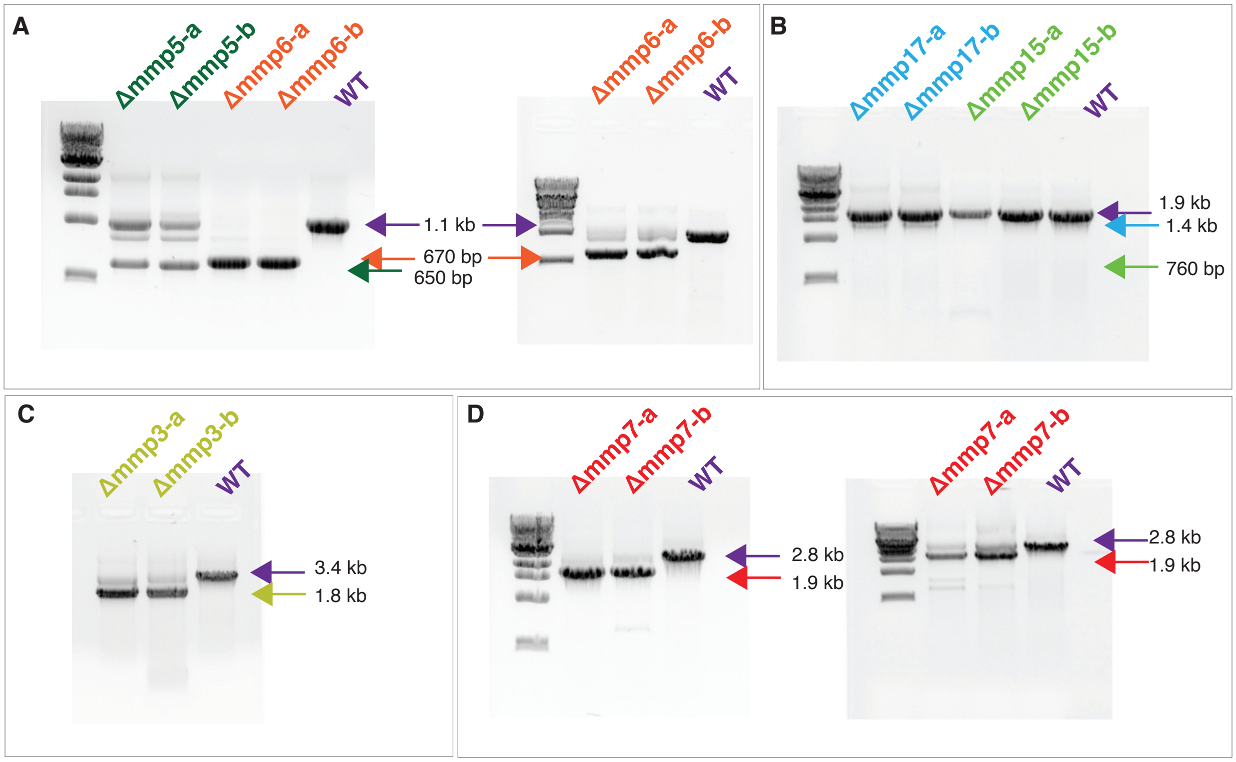


**Fig S3**. Attempts to delete genes in the MCR activation operon fail to yield clean deletion mutants. Agarose gels showing the PCR product generated with primers flanking the gene of interest from puromycin resistant colonies post transformation of the CRISPR-editing plasmid. (a) Diagnostic PCR for puromycin-resistant colonies (labeled a-b) when WWM60 was transformed with CRISPR-editing plasmids to delete Mmp5 and Mmp6, respectively. The PCR product corresponding to the wildtype (WT) allele is shown in purple, the PCR product corresponding to the deletion of Mmp5 is shown in dark green, and the PCR product corresponding to the deletion of Mmp6 is shown in orange (left). The gel on the right shows the appearance of a PCR product corresponding to the WT allele after several passages of the Δmmp6 cultures. (b) Diagnostic PCR for puromycin-resistant colonies (labeled a-b) when WWM60 was transformed with CRISPR-editing plasmids to delete Mmp17 and Mmp15, respectively. The PCR product corresponding to the wildtype (WT) allele is shown in purple, the PCR product corresponding to the deletion of Mmp17 is shown in light blue, and the PCR product corresponding to the deletion of Mmp15 is shown in light green. (c) Diagnostic PCR for puromycin-resistant colonies (labeled a-b) when WWM60 was transformed with CRISPR-editing plasmids to delete Mmp3. The PCR product corresponding to the wildtype (WT) allele is shown in purple and the PCR product corresponding to the deletion of Mmp3 is shown in chartreuse. (d) Diagnostic PCR for puromycin-resistant colonies (labeled a-b) when WWM60 was transformed with CRISPR-editing plasmids to delete Mmp7. The PCR product corresponding to the wildtype (WT) allele is shown in purple and the

PCR product corresponding to the deletion of Mmp3 is shown in red (left). The gel on the right shows the increased appearance of the PCR product corresponding to the WT allele after several round of passages of the Δmmp7 cultures.

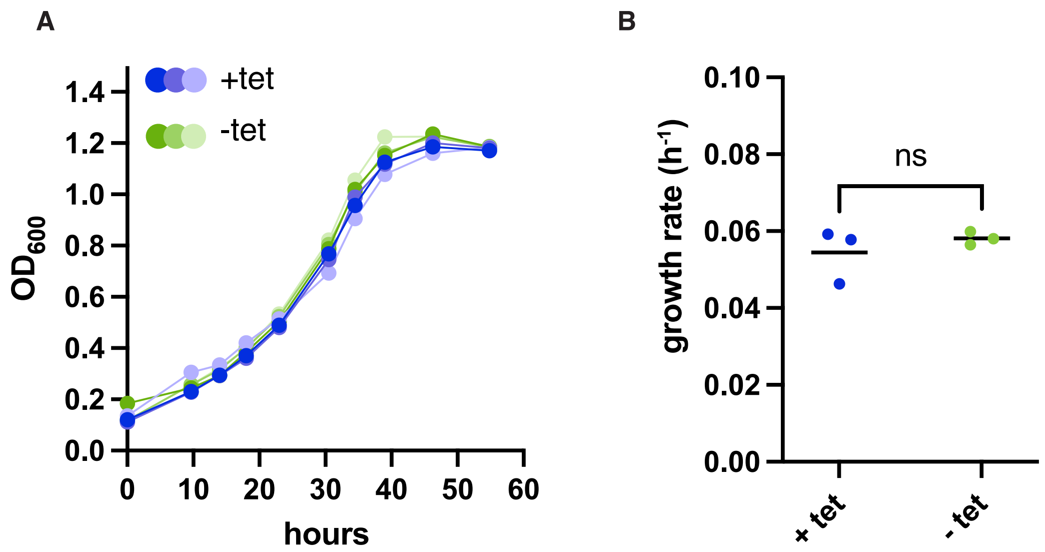


**Fig S4**. Analysis of growth when component A2 is overexpressed in *M. acetivorans*. (a) Growth curves of DDN122 expressing component A2 from the P*mcrB*(tetO1) tetracycline inducible promoter when expression of component A2 is induced by addition of 100 µg/mL tetracycline (+ tet, blue dots) or uninduced in the absence of tetraclycine (- tet, green dots). Cells were grown in 50 mM TMA-HS media supplemented with 2 µg/mL puromycin and 100 µg/mL tetracycline, if protein expression was induced (blue dots). Three replicates growth curves were conducted for each condition. (b) Growth rates corresponding to the curves plotted in (a). Statistical significance was tested by unpaired Welch's t test. ns indicates a p-value > 0.05.

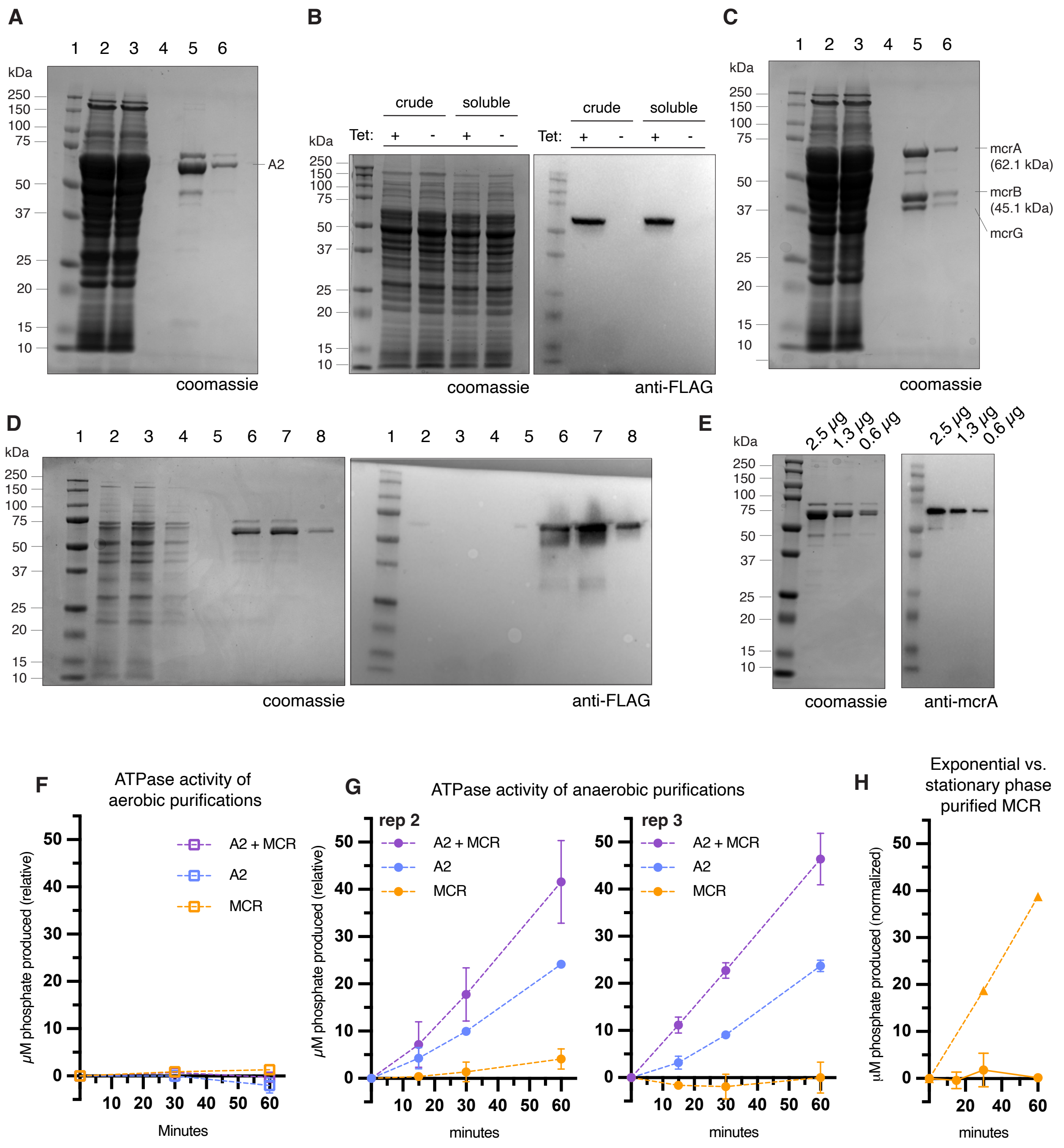


**Fig S5.** Purification of component A2 and MCR to determine ATPase activity. (a) SDS-PAGE gel representing various steps in the anaerobic affinity purification of component A2 using a Streptactin resin. Lanes as indicated are (1) ladder, (2) cleared lysate, (3) flow through, (4) second wash, (5) first elution, (6) second elution. (b) anti-FLAG western blot (right) showing the inducible production of component A2 in crude and soluble cell lysates of DDN122 and the corresponding SDS-PAGE gel (right) as a protein loading control. (c) SDS-PAGE Coomassie-stained gel showing the purification of MCR with the same lanes as (a). (d) anti-FLAG western blot (right) and SDS-PAGE Coomassie stained gel (left) as a protein loading control of component A2 purification with lanes as follows: (1) ladder, (2) cell lysate, (3) flow through, (4) first wash, (5) second wash, (6) first elution, (7) second elution, and (8) third elution. (e) anti-mcrA western blot (right) and SDS-PAGE Coomassie-stained gel (left), as a protein loading control, of purified component A2 diluted 2X from 2.5 μg to 0.6 μg per lane as indicated. (f) Aerobic ATPase assay of component A2 alone (blue), MCR alone (purple), and component A2 combined with MCR (orange). Each protein was purified under aerobic conditions. Each reaction contained 500 μg/mL of each protein in buffer with 200 μM ATP, 10 mM $MgCl_2$, 20 mM HEPES, 300 mM NaCl, and 1% glycerol. Reactions were incubated at 37 °C. Inorganic phosphate production was measured at 0, 30, and 60 minutes using malachite green reagent. (g) Additional biological replicates of anaerobic ATPase activity with two additional independent purification of component A2. Each reaction contained 500 μg/mL of each protein indicated with 200 μM ATP, 10 mM $MgCl_2$, 20 mM HEPES, 300 mM NaCl, and 1% glycerol. Reactions were incubated at 37 °C inside the anaerobic chamber. Inorganic phosphate production was measured at 0, 15, 30 and 60 minutes using malachite green reagent. (h) ATPase activity of MCR purified from exponential phase cells (triangles) with the same reaction conditions as (f), except time points were taken at 0, 30 and 60 minutes. The data from figure 1h for stationary phase purified MCR is included (circles) is included for comparison.

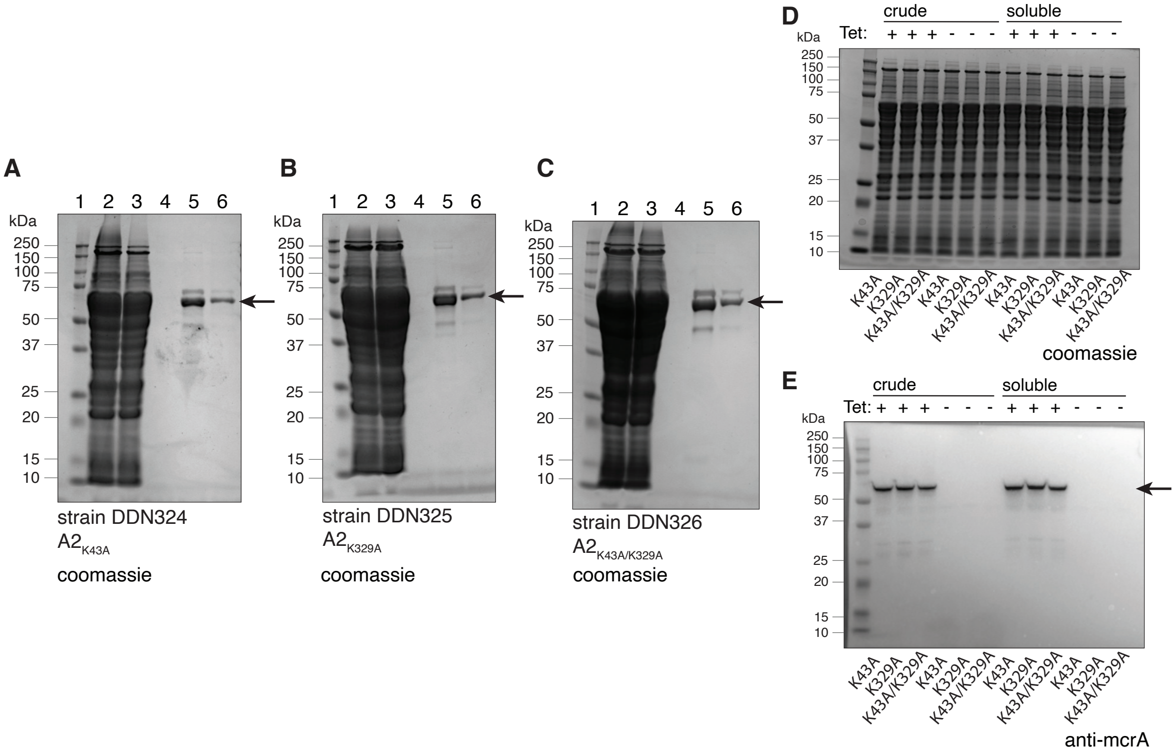


**Fig S6.** Purification of lysine point mutants of component A2. (a-c) SDS-PAGE Coomassie-stained gel showing the purification of (a) K43A, (b) K329A, and (c) K43A/K329A variants of component A2. Lanes as indicated are (1) ladder, (2) clear lysate, (3) flow through, (4) second wash, (5) first elution, (6) second elution. (d-e) anti-FLAG Western blot (e) showing the inducible production of each lysine point mutant in crude and soluble cell lysate and (d) the corresponding SDS-PAGE Coomassie-stained gel as a protein loading control.

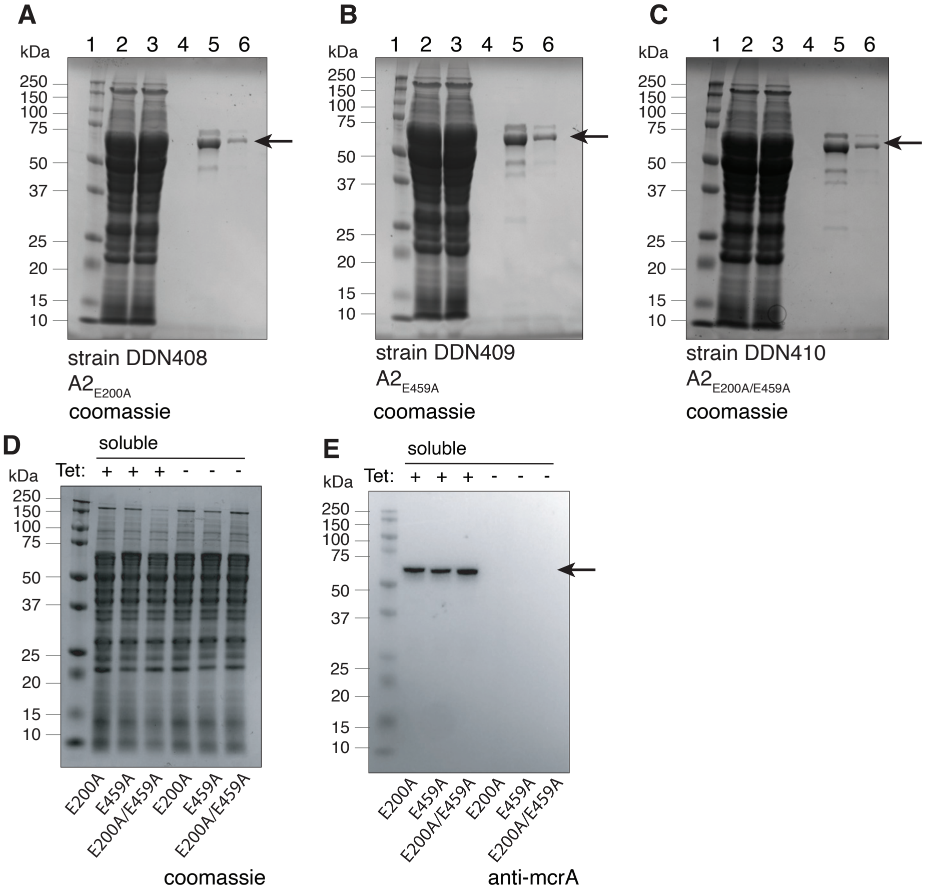


**Fig S7.** Purification of glutamate point mutants of component A2. (a-c) SDS-PAGE Coomassie-stained gel showing the purification of (a) E200A, (b) E459A, and (c) E200A/E459A. Lanes as indicated are (1) ladder, (2) clear lysate, (3) flow through, (4) second wash, (5) first elution, (6) second elution. (d-e) anti-FLAG Western blot (e) showing the inducible production of each glutamate point mutant in soluble cell lysates and the corresponding (d) SDS-PAGE Coomassie-stained gel as a protein loading control.

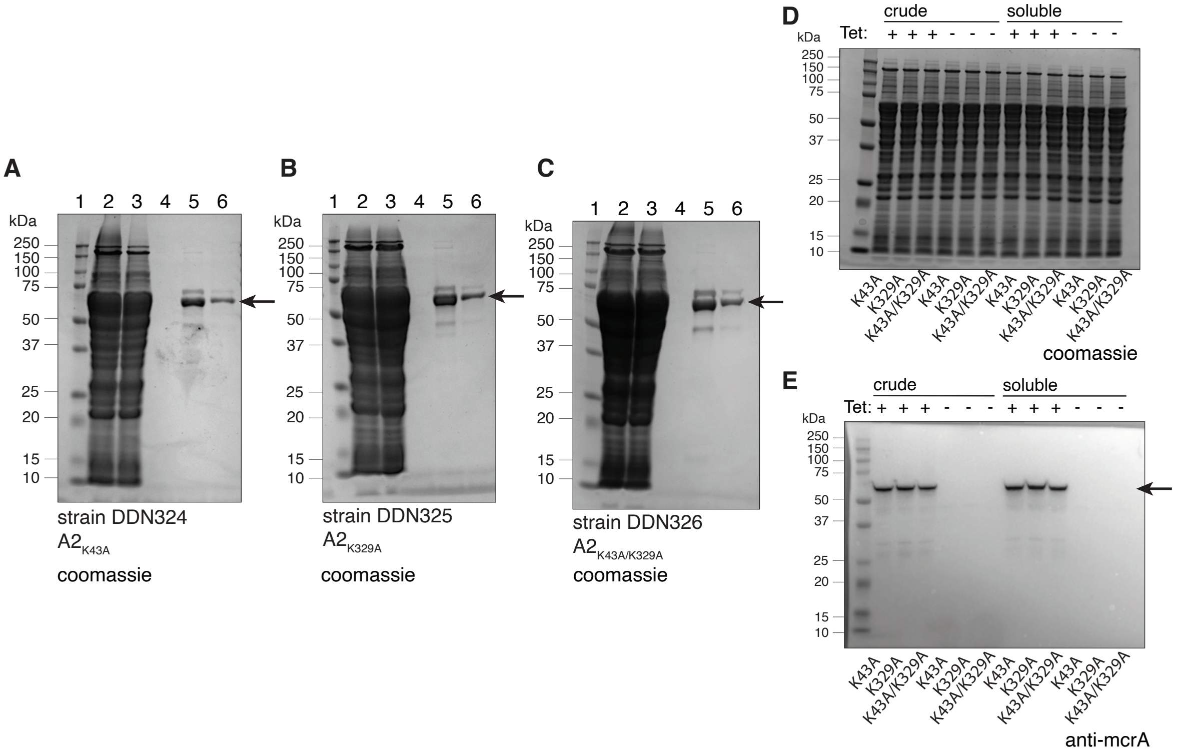


**Fig S8.** Purification of cysteine point mutants of component A2. (a) SDS-PAGE Coomassie-stained gel showing the purification of C70A/C73A/C86A/C89A point mutant of component A2. Lanes as indicated are (1) ladder, (2) clear lysate, (3) flow through, (4) second wash, (5) first elution, (6) second elution. (b-c) anti-FLAG Western blot (c) showing the inducible production of the cysteine point mutant in soluble cell lysates and the corresponding (b) SDS-PAGE Coomassie-stained gel as a protein loading control.

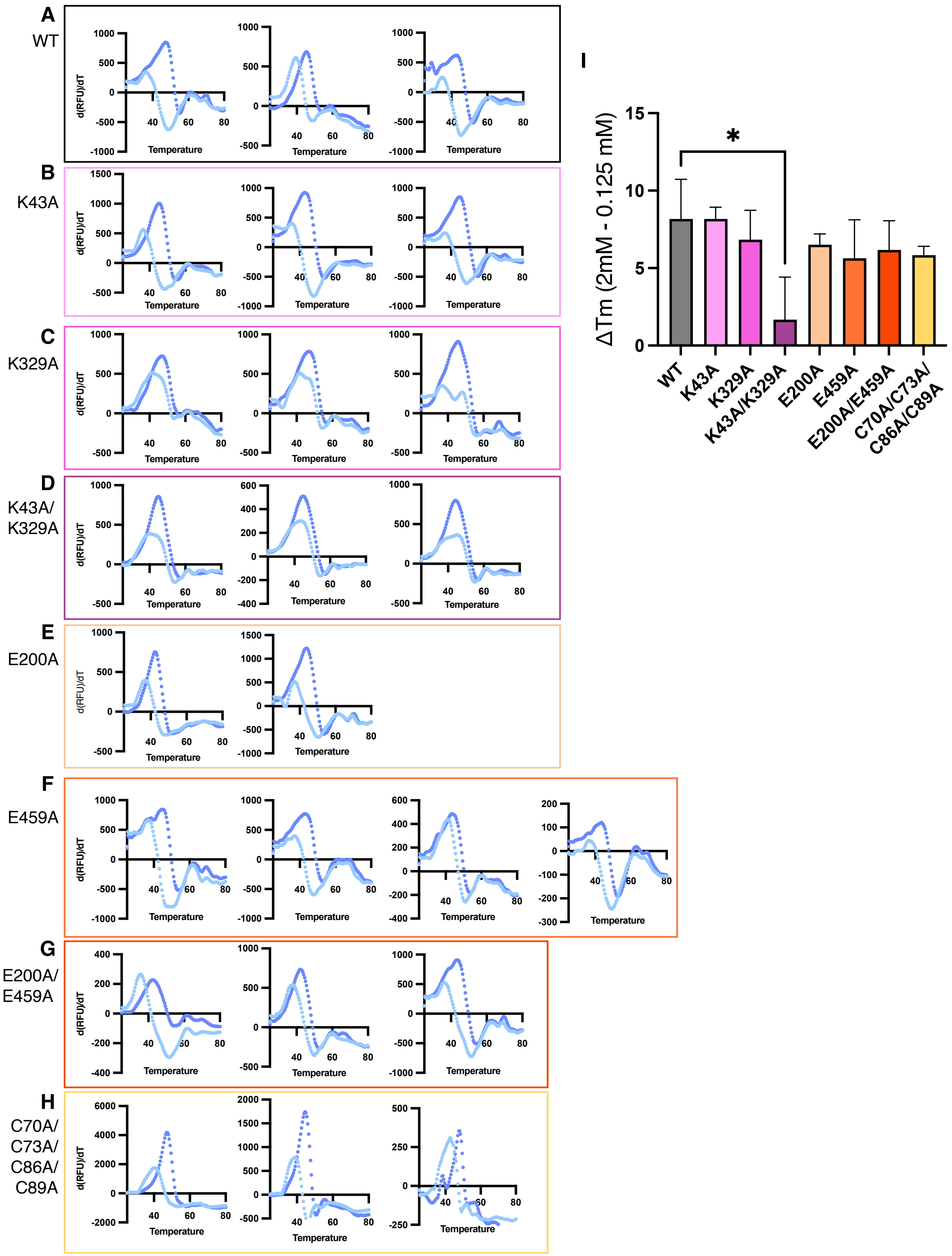

A
WT
B
K43A
C
K329A
D
K43A/
K329A
E
E200A
F
E459A
G
E200A/
E459A
H
C70A/
C73A/
C86A/
C89A
d(RFU)/dT
Temperature
I
ΔTm (2mM - 0.125 mM)
*
WT
K43A
K329A
K43A/K329A
E200A
E459A
E200A/E459A
C70A/C73A/
C86A/C89A

**Fig S9**. Change in melting temperature (Tm) with low (0.125 mM) and high (2 mM) concentrations of ATP across all variants of component A2. (a-h) Differential scanning fluorimetry of independent biological replicates of each point mutant and the wildtype (WT) allele (as indicated) with 0.125 mM (light blue) and 2 mM (dark blue) ATP. (i) summary of the changes in Tm found across panels (a-h) colored according to each point mutant in panels a-h. Error bars represent standard deviation of three replicates (except for E200A and E459A, which represent the data for two and four protein preparations, respectively) and Welch's unpaired t-test was performed to assess significance; * indicates P-value < 0.05.

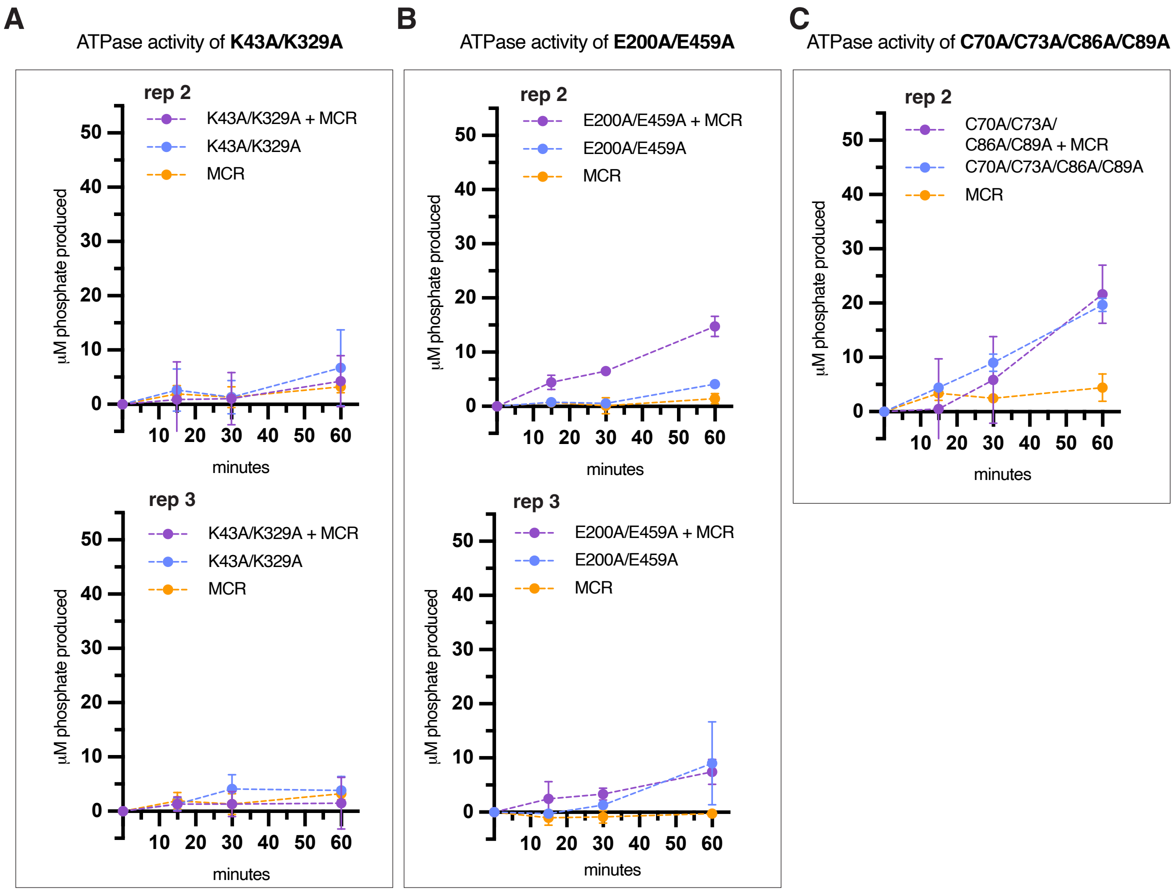


**Fig S10.** Independent replicates of the ATPase assay for K43A/K329A, E200A/K459A, and C70A/C73A/C86A/C89A mutants of Component A2. (a-c) Replicate anaerobic ATPase assays performed using independent purifications of each component A2 mutant. Each reaction contained 500 µg/mL of each protein indicated with 200 µM ATP, 10 mM $MgCl_2$, 20 mM HEPES, 300 mM NaCl, and 1% glycerol. Reactions were incubated at 37 °C. Inorganic phosphate production was measured at 0, 15, 30, and 60 minutes using malachite green reagent. Note: Replicate 1 for each mutant is shown in main text Figure 3.

**A** ATPase activity of K43A

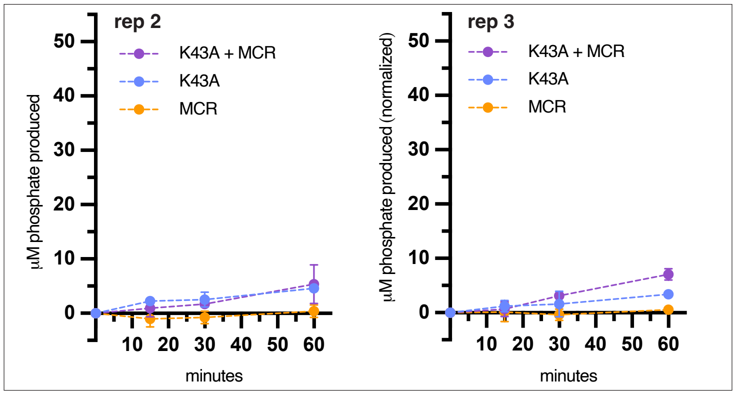


**B** ATPase activity of K329A

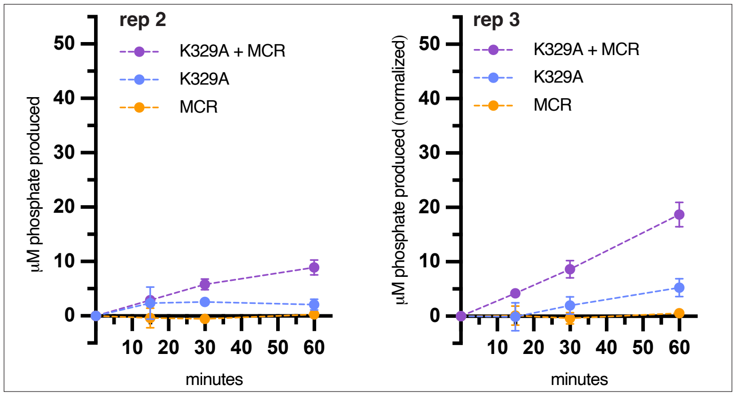


**C** ATPase activity of E200A

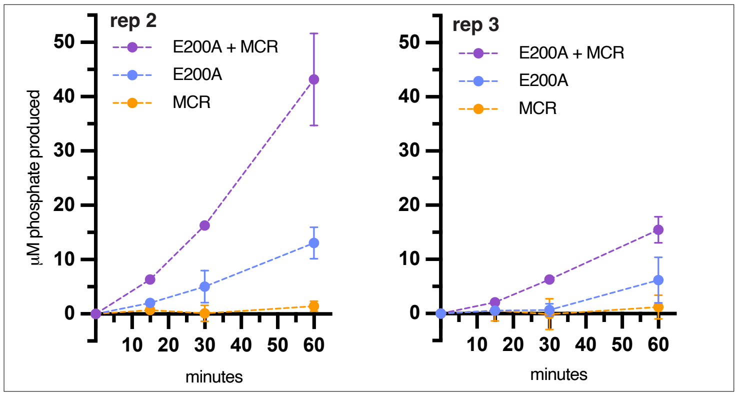


**D** ATPase activity of E459A

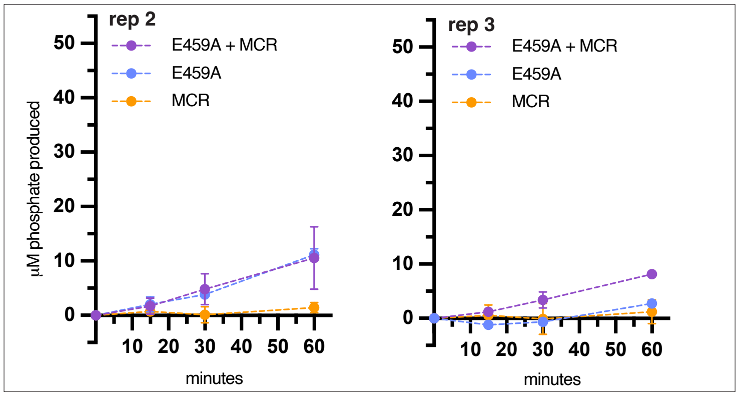

**Fig S11.** Independent replicates of the ATPase assay for K43A, K329A, E200A, and E459A mutants of component A2. (a-d) Replicate anaerobic ATPase assays performed using independent purifications of each component A2 mutant. Each reaction contained 500 µg/mL of each protein indicated with 200 µM ATP, 10 mM $MgCl_2$, 20 mM HEPES, 300 mM NaCl, and 1% glycerol. Reactions were incubated at 37 °C. Inorganic phosphate production was measured at 0, 15, 30, and 60 minutes using malachite green reagent. The same MCR fraction was used between K43A rep 3 and K329A rep 3, E200A rep 2 and E459A rep 2, and E200A rep 3 and E459A rep 3, so the MCR alone data is plotted across both plots in these panels for comparison. Note: Replicate 1 for each mutant is shown in main text Figure 3.

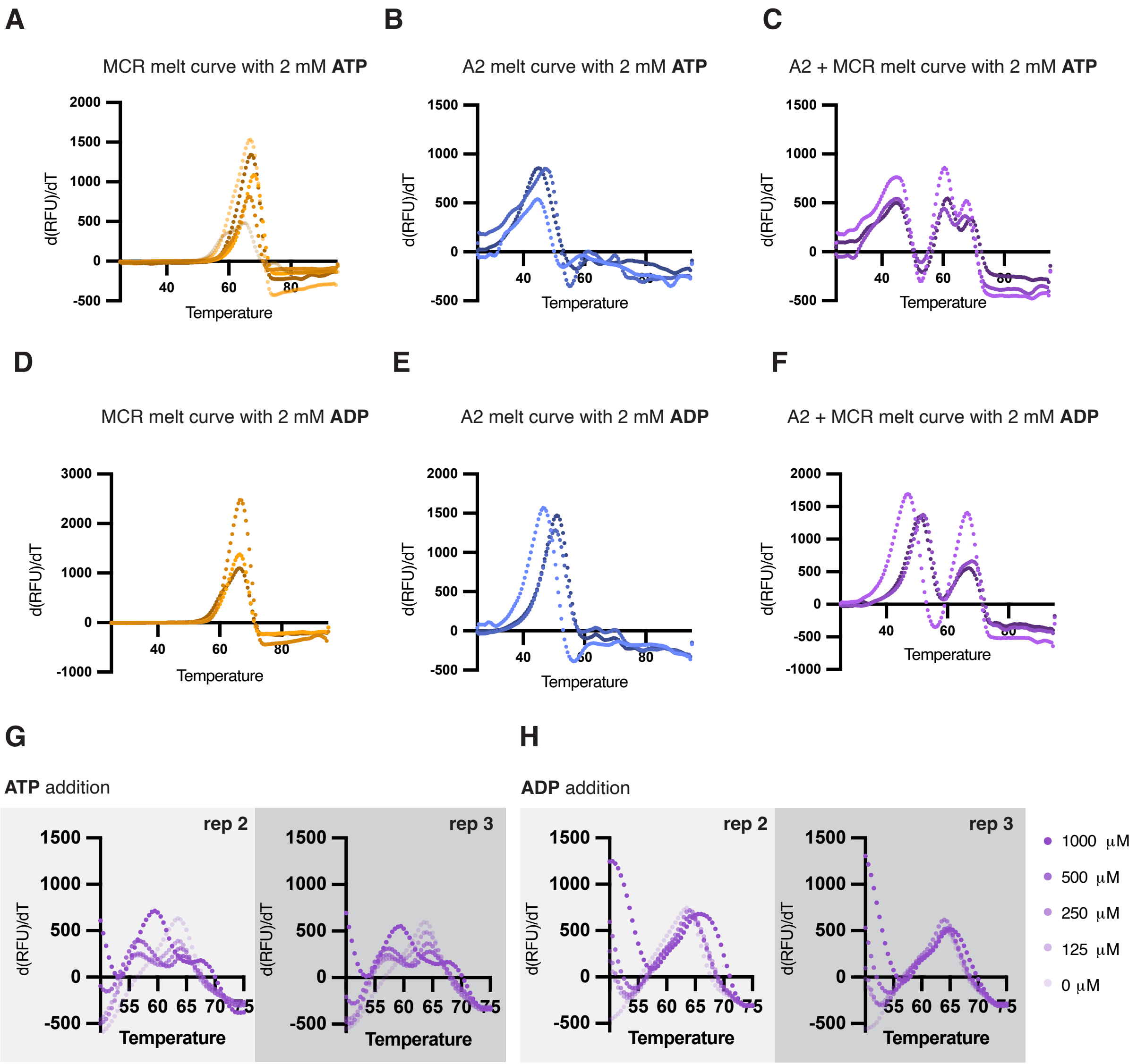


**Fig S12.** Differential Scanning Fluorimetry (DSF) of component A2 and MCR with ATP or ADP. (a-c) Multiple independent purifications of MCR (a), component A2 (b), or MCR + component A2 (c). Samples contained 0.5 mg/mL each protein, 2 mM ATP, 10 mM $MgCl_2$, 20 mM HEPES, 300 mM NaCl, and 1% glycerol. (d-f) Three independent purifications of MCR (d), component A2 (e), or MCR + component A2 (f). Samples contained 2 mM ADP, 10 mM $MgCl_2$, 20 mM HEPES, 300 mM NaCl, and 1% glycerol. (g-h) Two additional replicates of DSF melt curves with increasing concentrations of ATP (g) or ADP (h) and the same conditions as (a-f). Note: Replicate 1 is shown in main text Figure 4.

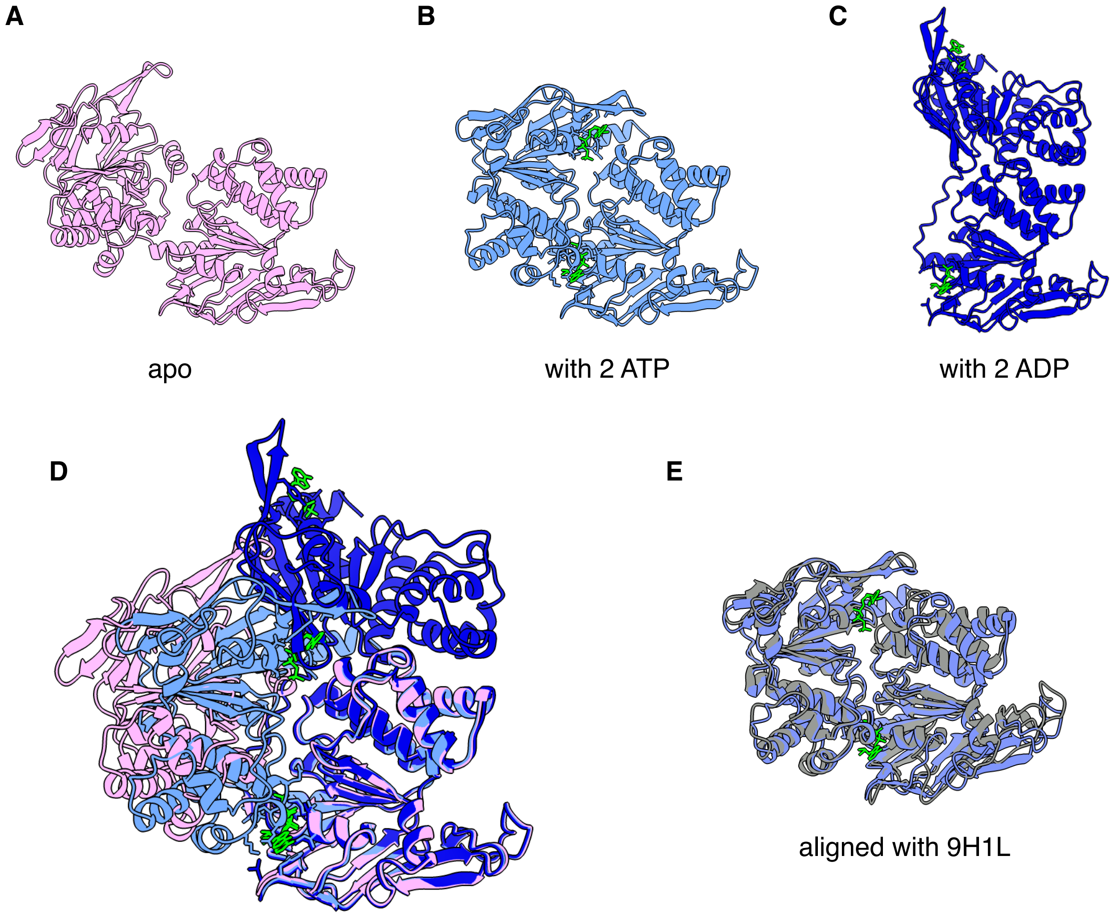


**Fig S13.** AlphaFold predicted structures of component A2 with and without nucleotides bound. (a) AlphaFold predicted structure of *M. acetivorans* component A2 without any nucleotide (pink). (b) AlphaFold predicted structure of *M. acetivorans* component A2 with 2 molecules of ATP (light blue). (c) AlphaFold predicted structure of *M. acetivorans* component A2 with 2 molecules of ADP (dark blue). (d) alignment of the predicted structures in a-c maintaining the same orientation with apo-protein (pink), ATP-bound (light blue), and ADP-bound (dark blue) forms. RMSD of apo component A2 aligned with ATP-bound component A2 is 10.262 Å, RMSD of apo component A2 aligned with ADP-bound component A2 is 30.107 Å. (e) AlphaFold predicted structure of *M. acetivorans* component A2 with 2 molecules of ATP (light blue) aligned with the resolved A2 structure from PDB: 9H1L (grey). RMSD is 2.667 Å. Each panel shows the relevant nucleotide in green.

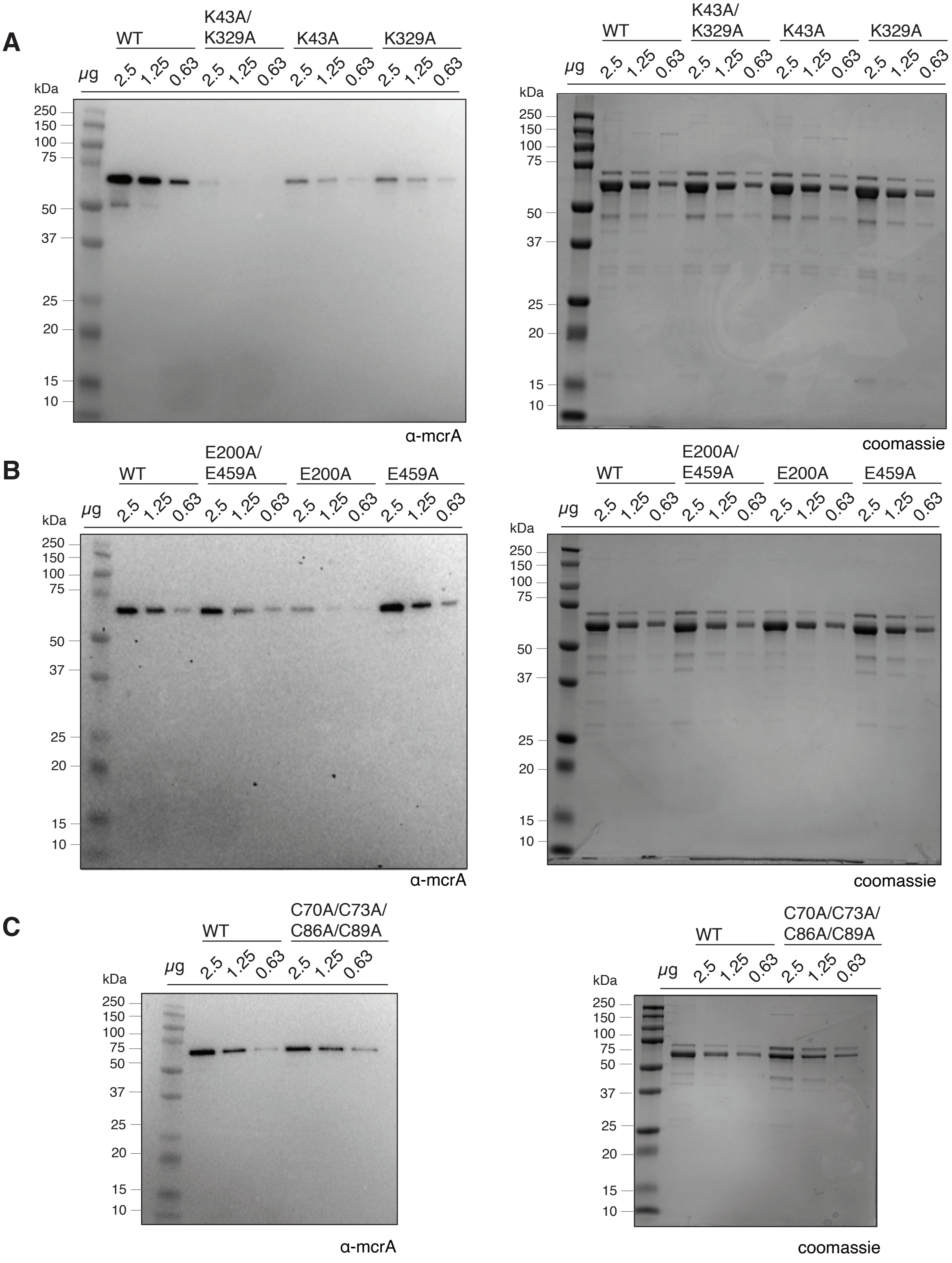
A
WT
K43A/
K329A
K43A
K329A
μg
2.5
1.25
0.63
kDa
250
150
100
75
50
37
25
20
15
10
α-mcrA
coomassie
B
WT
E200A/
E459A
E200A
E459A
C
WT
C70A/C73A/
C86A/C89A

**Fig S14.** Full images of western blots shown in Figure 5a. (a-b) Uncropped western blots (left) with anti-McrA specific antibodies and Coomassie stained SDS-PAGE gels (right) as a protein loading control for (a) K/A mutants, (b) E/A mutants, and (c) C/A mutant as indicated.

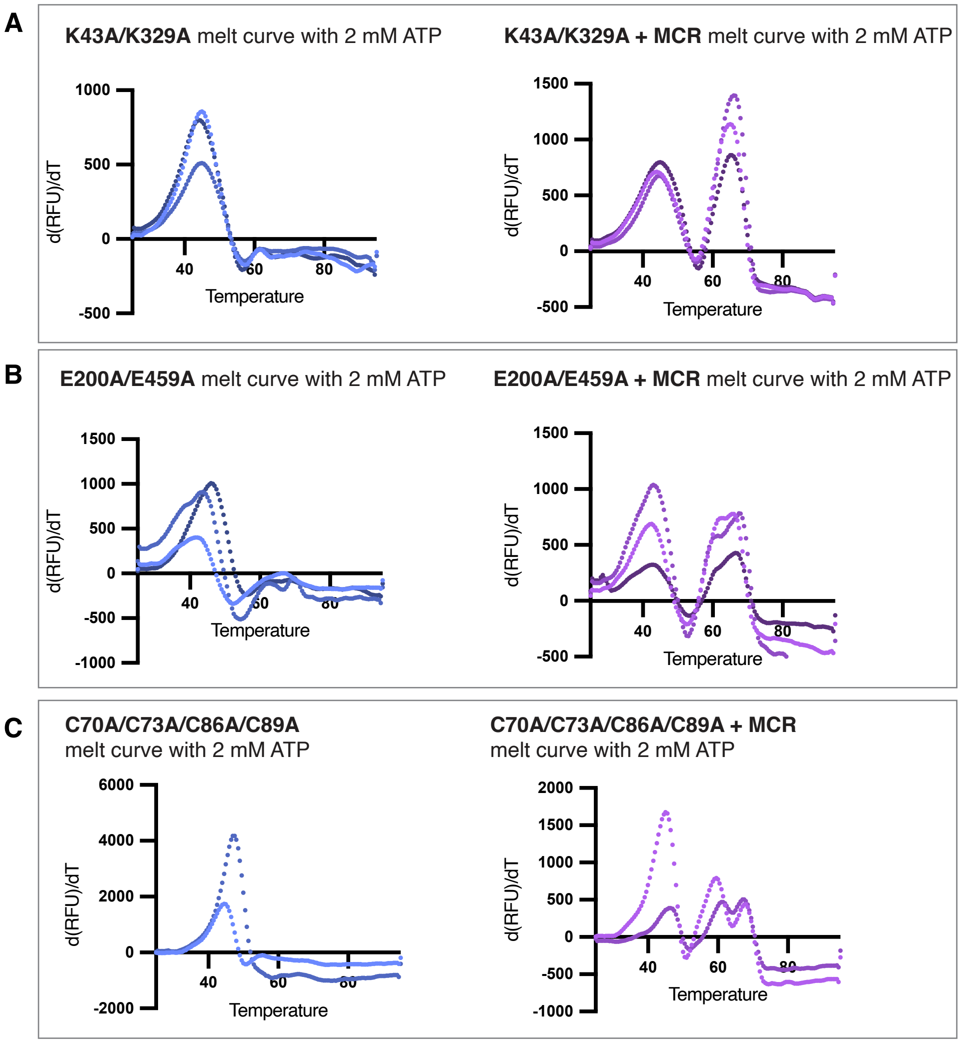


**Fig S15.** Differential scanning fluorimetry of K43A/K329A, E200A/E459A, and C70A/C73A/C86A/C89A mutants of component A2 + MCR in the presence of ATP. (a-b) Three independent purifications of (a) K43A/K329A component A2, or (b) MCR + K43A/K329A component A2. Samples contained 2 mM ATP, 10 mM $MgCl_2$, 20 mM HEPES, 300 mM NaCl, and 1% glycerol. (c-d) Three independent purifications of (c) E200A/E459A component A2, or (d) MCR + E200A/E459A component A2, with the same sample conditions as (a). (e-f) Two independent purifications of (e) C70A/C73A/C86A/C89A component A2, or (f) MCR + C70A/C73A/C86A/C89A component A2 (f), with the same sample conditions as (a). All proteins are at a concentration of 0.5 mg/mL.

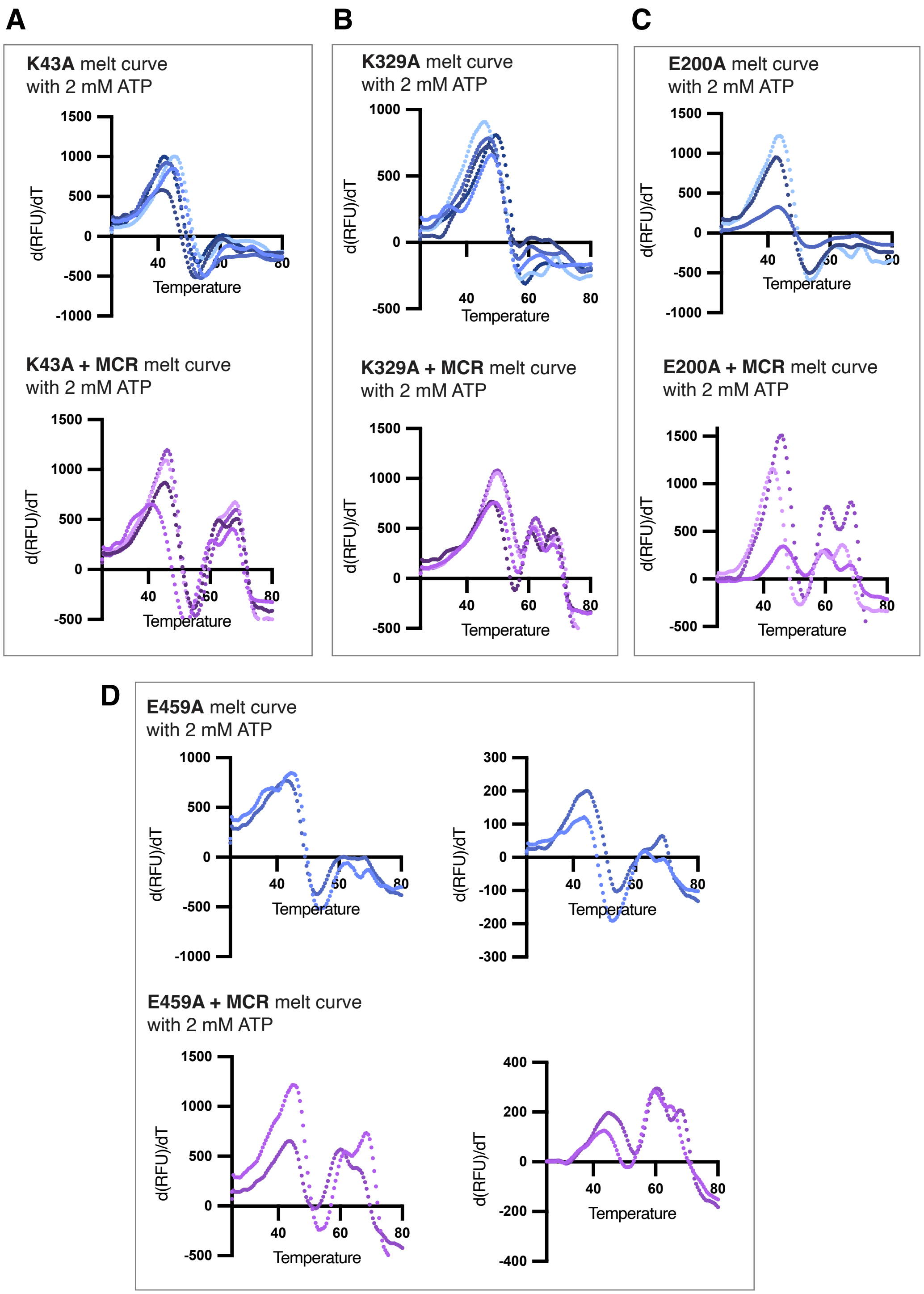


**Fig S16.** Differential Scanning Fluorimetry of K43A, K329A, E200A, and E459A mutants of component A2 + MCR with ATP. (a) Independent purifications of K43A (top), or MCR + K43A (bottom). Samples contained 2 mM ATP, 10 mM $MgCl_2$, 20 mM HEPES, 300 mM NaCl, and 1% glycerol. (b) Independent purifications of K329A (top), or MCR + K329A (bottom)., with the same sample conditions as (a). (c) Independent purifications of E200A (top), or MCR + E200A (bottom), with the same sample conditions as (a). (d) Independent purifications of E459A (top), or MCR + E459A (bottom), with the same sample conditions as (a). All proteins are at a concentration of 0.5 mg/mL.

**Table S1.** Mutations predicted in DDN122 by whole-genome sequencing. Mutations were predicted using Breseq version v0.35.5 using default parameters.

| Position | Mutation | Annotation | Location | Present in WWM73 |
|---|---|---|---|---|
| 487,691 | +C | intergenic (+91/-1171) | *MA_0404* → / → *isf-3* | yes |
| 954,776 | $(A)_{5\rightarrow 6}$ | coding (27/765 nt) | *MA_0806* ← | yes |
| 1,327,728 | Δ1 bp | intergenic (+539/+2434) | *MA_1110* → / ← *MA_1114* | yes |
| 2,100,489 | 2 bp→CT | intergenic (-200/+180) | *MA_1738* ← / ← *MA_1739* | yes |
| 2,100,494 | G→T | intergenic (-205/+176) | *MA_1738* ← / ← *MA_1739* | yes |
| 2,548,151 | +C | coding (6598/6654 nt) | *MA_2045* ← | yes |
| 2,850,254 | A→G | F64L (TTC→CTC) | *serC* ← | yes |
| 2,880,667 | T→G | H313Q (CAT→CAG) | *MA_2328* → | yes |
| 3,446,809 | Δ1 bp | coding (771/774 nt) | *MA_2751* → | yes |
| 4,309,060 | Δ1 bp | coding (108/12681 nt) | *MA_3471* ← | yes |
| 4,888,175 | Δ1 bp | intergenic (-67/-228) | *MA_3972* ← / → *MA_3973* | yes |
| 4,958,953 | +C | intergenic (+2396/-395) | *MA_4028* → / → *MA_4030* | yes |

**Table S2.** Identification of individual SDS-PAGE bands of in-gel digests from component A2 preparation showing tryptic peptides and resulting proteins identified.

| **~70 kDa band** | | | | |
|---|---|---|---|---|
| **Glutamyl-tRNA(Gln) amidotransferase subunit E, MA_2862, MW: 70.8** | | | | |
| **Protein score CI% 100%** | | | | |
| Calc. Mass | Observ. Mass | Sequence | Ion score | C. I. % |
| 841.4638 | 841.4547 | EVQPGRR | | |
| 852.4396 | 852.4212 | EKLFCR | | |
| 924.4421 | 924.4196 | LGTEFSDR | | |
| 998.5992 | 998.5614 | LAIEAVINR | | |
| 998.5992 | 998.5614 | LAIEAVINR | 52 | 99.977 |
| 1013.6716 | 1013.6249 | GSILAALLKK | | |
| 1015.553 | 1015.5273 | QDVNISIAR | | |
| 1016.6251 | 1016.5781 | QLNLLFIR | 49 | 99.95 |
| 1016.6251 | 1016.5781 | QLNLLFIR | | |
| 1080.5432 | 1080.5029 | RLGTEFSDR | | |
| 1096.6725 | 1096.627 | TLVGIVPEIR | 62 | 99.998 |
| 1096.6725 | 1096.627 | TLVGIVPEIR | | |
| 1108.5746 | 1108.54 | RFASESGLNK | | |
| 1124.5582 | 1124.525 | GDSIVYSLDR | | |
| 1173.6222 | 1173.5884 | DANVNSTLIAR | | |
| 1279.6715 | 1279.6223 | EVAEYIGMVLR | 94 | 100 |
| 1279.6715 | 1279.6223 | EVAEYIGMVLR | | |
| 1295.6664 | 1295.6144 | EVAEYIGMVLR | | |
| 1347.5852 | 1347.5331 | DIGDSDFEFFR | | |
| 1347.5852 | 1347.5331 | DIGDSDFEFFR | 103 | 100 |
| 1361.7423 | 1361.6803 | GLFAAISNQEIAK | | |
| 1371.7114 | 1371.6622 | EAIEGIPEETRK | | |
| 1379.6914 | 1379.6462 | IVVDGSNTSGFQR | | |
| 1440.8057 | 1440.7471 | GVQALDLIEDIVR | 81 | 100 |
| 1440.8057 | 1440.7471 | GVQALDLIEDIVR | | |
| 1475.7086 | 1475.6986 | MEKYDYSELGLK | | |
| 1507.7863 | 1507.736 | KIVVDGSNTSGFQR | | |
| 1577.9261 | 1577.863 | LGIPLVEIATAPDIR | 61 | 99.997 |
| 1577.9261 | 1577.863 | LGIPLVEIATAPDIR | | |
| 1596.9067 | 1596.8505 | GVQALDLIEDIVRR | | |
| 1623.8224 | 1623.776 | IEEKGDSIVYSLDR | | |
| 1716.8075 | 1716.7592 | TAFLASDGYIETSEGR | 78 | 100 |

| | | | | |
|---|---|---|---|---|
| 1716.8075 | 1716.7592 | TAFLASDGYIETSEGR | | |
| 1745.8649 | 1745.8066 | LFNMKPVDQMHVMR | | |
| 1745.8649 | 1745.8066 | LFNMKPVDQMHVMR | 78 | 100 |
| 1761.8597 | 1761.803 | LFNMKPVDQMHVMR | | |
| 1807.9113 | 1807.8192 | LGLSAFDPEEVENFIK | | |
| 1870.9592 | 1870.9004 | ALPDGNTAYMRPLPGAAR | 48 | 99.945 |
| 1870.9592 | 1870.9004 | ALPDGNTAYMRPLPGAAR | | |
| 1886.9541 | 1886.8969 | ALPDGNTAYMRPLPGAAR | | |
| 1922.8436 | 1922.7898 | AYDTTCLIENDEEPPR | | |
| 2288.0862 | 2288.0342 | DAIGAGPEDAFVMVADEPEKAR | | |
| 2407.1082 | 2407.0491 | AYDTTCLIENDEEPPREL | | |

**~60 kDa band**

**ABC transporter, ATP-binding protein, MA_3998, MW: 59.6**

**Protein score CI% 100%**

| Calc. Mass | Observ. Mass | Sequence | Ion score | C. I. % |
|---|---|---|---|---|
| 844.5073 | 844.4847 | IAIMLQR | | |
| 860.5022 | 860.4769 | IAIMLQR | | |
| 908.4948 | 908.4698 | RYISVDR | | |
| 968.6502 | 968.611 | IALAQVLIK | | |
| 976.5461 | 976.5302 | LSLYDPIR | | |
| 982.5866 | 982.5566 | TILMHVLR | 57 | 99.994 |
| 982.5866 | 982.5566 | TILMHVLR | | |
| 998.5815 | 998.5537 | TILMHVLR | | |
| 1000.6084 | 1000.5824 | RIAIMLQR | | |
| 1003.5166 | 1003.5363 | ELSGGEKQR | | |
| 1016.6033 | 1016.5746 | RIAIMLQR | | |
| 1071.5105 | 1071.4856 | TFALYGDER | 66 | 100 |
| 1071.5105 | 1071.4856 | TFALYGDER | | |
| 1104.6411 | 1104.6071 | LSLYDPIRK | | |
| 1109.6677 | 1109.6251 | NVLVGEPIIR | | |
| 1109.6677 | 1109.6251 | NVLVGEPIIR | 31 | 97.283 |
| 1121.5837 | 1121.5616 | NLTVDFDGLK | | |
| 1179.535 | 1179.5647 | VGDEWVDMTK | | |
| 1184.4558 | 1184.4395 | MTDEMSEGER | | |
| 1206.7205 | 1206.6803 | LVHEAIINAVK | | |
| 1350.8467 | 1350.814 | IALAQVLIKEPR | | |
| 1350.8467 | 1350.814 | IALAQVLIKEPR | 66 | 100 |
| 1493.6107 | 1493.7108 | MTDEMSEGERHR | | |
| 1508.8066 | 1508.765 | AVELLEEVNLSHR | 88 | 100 |

| | | | | |
|---|---|---|---|---|
| 1508.8066 | 1508.765 | AVELLEEVNLSHR | | |
| 1636.9016 | 1636.8674 | KAVELLEEVNLSHR | 77 | 100 |
| 1636.9016 | 1636.8674 | KAVELLEEVNLSHR | | |
| 1710.9458 | 1710.9009 | ILMGILPPTSGEVEVR | | |
| 1710.9458 | 1710.9009 | ILMGILPPTSGEVEVR | 95 | 100 |
| 1726.9408 | 1726.902 | ILMGILPPTSGEVEVR | | |
| 2021.9927 | 2021.9484 | GTESFENISGEVIYHLAR | 160 | 100 |
| 2021.9927 | 2021.9484 | GTESFENISGEVIYHLAR | | |
| 2110.1101 | 2110.1047 | NPLLLLADEPTGTLDPMTAK | | |
| 2110.1501 | 2110.1047 | IVEIGRPGTVLAQLTEEER | 77 | 100 |

**~50 kDa band**

| **ABC transporter, ATP-binding protein, MA_3998, MW: 59.6** | | | | |
|---|---|---|---|---|
| **Protein score CI% 100%** | | | | |
| Calc. Mass | Observ. Mass | Sequence | Ion score | C. I. % |
| 844.5073 | 844.5098 | IAIMLQR | | |
| 908.4948 | 908.4931 | RYISVDR | | |
| 976.5461 | 976.5475 | LSLYDPIR | | |
| 982.5866 | 982.5903 | TILMHVLR | | |
| 982.5866 | 982.5903 | TILMHVLR | 42 | 99.755 |
| 998.5815 | 998.5963 | TILMHVLR | | |
| 1000.6084 | 1000.6046 | RIAIMLQR | | |
| 1003.5166 | 1003.5557 | ELSGGEKQR | | |
| 1016.6033 | 1016.622 | RIAIMLQR | | |
| 1071.5105 | 1071.5198 | TFALYGDER | 38 | 99.423 |
| 1071.5105 | 1071.5198 | TFALYGDER | | |
| 1104.6411 | 1104.6388 | LSLYDPIRK | | |
| 1109.6677 | 1109.6621 | NVLVGEPIIR | | |
| 1109.6677 | 1109.6621 | NVLVGEPIIR | 43 | 99.825 |
| 1179.535 | 1179.6029 | VGDEWVDMTK | | |
| 1206.7205 | 1206.7023 | LVHEAIINAVK | | |
| 1493.6107 | 1493.7355 | MTDEMSEGERHR | | |
| 1508.8066 | 1508.7999 | AVELLEEVNLSHR | | |
| 1508.8066 | 1508.7999 | AVELLEEVNLSHR | 73 | 100 |
| 1636.9016 | 1636.897 | KAVELLEEVNLSHR | | |
| 1710.9458 | 1710.9396 | ILMGILPPTSGEVEVR | | |
| 1710.9458 | 1710.9396 | ILMGILPPTSGEVEVR | 44 | 99.836 |
| 2021.9927 | 2021.9875 | GTESFENISGEVIYHLAR | | |
| 2110.1501 | 2110.1387 | IVEIGRPGTVLAQLTEEER | | |

**~45 kDa band**

| **Methyl coenzyme-M reductase subunit beta, MA_4550, MW: 45** | | | | |
|---|---|---|---|---|
| **Protein score CI% 100%** | | | | |
| Calc. Mass | Observ. Mass | Sequence | Ion Score | C. I. % |
| 858.5407 | 858.5487 | IILDTKR | | |
| 1071.6157 | 1071.6185 | SLLIQAPSSR | | |
| 1071.6157 | 1071.6185 | SLLIQAPSSR | 34 | 98.555 |
| 1131.5891 | 1131.5687 | AVEDGVISVDK | | |
| 1383.6387 | 1383.6803 | SDTVDIYDDRGK | | |
| 1411.7474 | 1411.7524 | NIMANHIAAITSR | | |
| 1411.7474 | 1411.7524 | NIMANHIAAITSR | 75 | 100 |
| 1416.7693 | 1416.7738 | DGTIGTVIESIVGR | | |
| 1427.7423 | 1427.7592 | NIMANHIAAITSR | | |
| 1658.8782 | 1658.879 | LLESNVDIMSLAPTR | 68 | 100 |
| 1658.8782 | 1658.879 | LLESNVDIMSLAPTR | | |

**~27 kDa band**

| **Methyl coenzyme-M reductase subunit gamma, MA_4547, MW: 27.6** | | | | |
|---|---|---|---|---|
| **Protein score CI% 100%** | | | | |
| Calc. Mass | Observ. Mass | Sequence | Ion Score | C. I. % |
| 1020.5333 | 1020.5558 | GATVHGHSVR | | |
| 1088.5524 | 1088.5701 | SYFAAINFR | | |
| 1088.5524 | 1088.5701 | SYFAAINFR | 62 | 99.997 |
| 1090.5813 | 1090.5734 | LEGGVIIMDK | | |
| 1109.4714 | 1109.6082 | ERDMEQCAK | | |
| 1368.6543 | 1368.6763 | DDAEVIEWVHR | | |
| 1554.7759 | 1554.7944 | EISDEDLTAVLGHR | | |
| 1568.7084 | 1568.7329 | LQEDGVMFDMLDR | | |
| 1724.8094 | 1724.8285 | LQEDGVMFDMLDRR | | |
| 1779.8075 | 1779.83 | VQMETEMTDPALAGMR | | |
| 1817.8452 | 1817.8657 | AYEAQYYPGATSVGANR | | |
| 1817.8452 | 1817.8657 | AYEAQYYPGATSVGANR | 64 | 99.999 |
| 1823.9609 | 1823.9769 | LREISDEDLTAVLGHR | | |
| 2140.9797 | 2141.0093 | YVQFADSMYNAPATPYF R | 81 | 100 |
| 2140.9797 | 2141.0093 | YVQFADSMYNAPATPYF | | |

| | | |
|---|---|---|
| | | R |
| 2170.0676 | 2170.0933 | VDNVAFRDDAEVIEWVHR |
| 2540.1545 | 2540.1855 | APGSDYPSTHPPLAEMGEPACSIR |

**~30 kDa band**

| **ABC transporter, ATP-binding protein, MA_3998, MW: 59.6** | | | | | |
|---|---|---|---|---|---|
| **Sequence coverage: 43%** | | | | | |
| Positions in Master Proteins | Theo. MH+ [Da] | DeltaM [ppm]: Mascot | Deltam/z [Da]: Mascot | Sequence | Confidence |
| 19918090 [415-424] | 1064.5371 | -1.32 | -0.0007 | AAGFEENKAK | High |
| 19918090 [430-441] | 1477.61584 | -3.32 | -0.00163 | MTDEMSEGERHR | High |
| 19918090 [430-441] | 1493.61075 | -0.29 | -0.00014 | MTDEMSEGERHR | High |
| 19918090 [9-18] | 1121.58372 | -0.37 | -0.00021 | NLTVDFDGLK | High |
| 19918090 [9-21] | 1433.79986 | -1.02 | -0.00073 | NLTVDFDGLKALK | High |
| 19918090 [278-287] | 1109.66772 | -0.06 | -0.00003 | NVLVGEPIIR | High |
| 19918090 [275-287] | 1494.86385 | -1.09 | -0.00081 | QEKNVLVGEPIIR | High |
| 19918090 [430-439] | 1216.44564 | 1.37 | 0.00083 | MTDEMSEGER | High |
| 19918090 [44-51] | 998.58155 | -1.24 | -0.00062 | TILMHVLR | High |
| 19918090 [470-478] | 944.57751 | -0.46 | -0.00022 | VAVTNSILK | High |
| 19918090 [351-360] | 1179.53505 | 0.75 | 0.00044 | VGDEWVDMTK | High |
| 19918090 [351-360] | 1195.52997 | 1.22 | 0.00073 | VGDEWVDMTK | High |
| 19918090 [124-132] | 1071.51055 | -0.23 | -0.00012 | TFALYGDER | High |
| 19918090 [430-439] | 1184.45581 | 0.86 | 0.00051 | MTDEMSEGER | High |

| | | | | | |
|---|---|---|---|---|---|
| 19918090 [454-469] | 1792.87071 | -2.49 | -0.00223 | IVIMDEPTGTMDPITK | High |
| 19918090 [305-314] | 1121.6201 | -0.92 | -0.00052 | AVDKISFDVK | High |
| 19918090 [155-167] | 1508.80673 | 1.59 | 0.0012 | AVELLEEVNLSHR | High |
| 19918090 [73-81] | 1109.5408 | 1.11 | 0.00062 | CGYIERPSK | High |
| 19918090 [315-329] | 1433.76347 | -2.04 | -0.00146 | EGEIFGLVGISGAGK | High |
| 19918090 [174-182] | 1003.5167 | -1.27 | -0.00063 | ELSGGEKQR | High |
| 19918090 [408-414] | 786.54475 | 0.43 | 0.00017 | KAIITLK | High |
| 19918090 [154-167] | 1636.9017 | 0.12 | 0.00006 | KAVELLEEVNLSHR | High |
| 19918090 [442-453] | 1350.84675 | 0.17 | 0.00011 | IALAQVLIKEPR | High |
| 19918090 [361-368] | 858.47919 | -0.73 | -0.00031 | LGVDNKGR | High |
| 19918090 [335-350] | 1710.94587 | -0.51 | -0.00044 | ILMGILPPTSGEVEVR | High |
| 19918090 [335-350] | 1726.94078 | -0.04 | -0.00003 | ILMGILPPTSGEVEVR | High |
| 19918090 [27-38] | 1227.69433 | -0.05 | -0.00003 | INEGEVVGILGK | High |
| 19918090 [212-222] | 1206.72048 | -0.59 | -0.00035 | LVHEAIINAVK | High |
| 19918090 [454-469] | 1760.88088 | -1.51 | -0.00133 | IVIMDEPTGTMDPITK | High |
| 19918090 [454-469] | 1776.8758 | -0.7 | -0.00062 | IVIMDEPTGTMDPITK | High |
| 19918090 [442-450] | 968.65028 | -0.59 | -0.00029 | IALAQVLIK | High |
| 19918090 [295-304] | 1163.65313 | -0.85 | -0.00049 | YISVDRGVVR | High |

**Methanogenesis Marker Protein 7, MA_3992, MW: 34.3**
**Sequence coverage: 37%**

| Positions in Master Proteins | Theo. MH+ [Da] | DeltaM [ppm]: Mascot | Deltam/z [Da]: Mascot | Sequence | Confidence |
|---|---|---|---|---|---|
| 499333395 [295-300] | 763.47125 | -0.18 | -0.00007 | NYILIK | High |
| 499333395 [218-230] | 1406.81142 | -0.88 | -0.00041 | QQAKDPLAVLPAR | High |
| 499333395 [117-126] | 1271.60486 | 0.61 | 0.00039 | RVSLSMDYER | High |
| 499333395 [172-187] | 1675.82859 | 0.55 | 0.00046 | TEDVPGADLYVGNIGR | High |
| 499333395 [195-210] | 1774.9069 | -1.38 | -0.00082 | TEEIEALDKLNEGVSK | High |
| 499333395 [245-259] | 1592.93702 | 0.79 | 0.00063 | VLSPAPLTLQLDGLR | High |
| 499333395 [231-244] | 1712.89998 | 1.83 | 0.00104 | VMKEIENQVPEIQR | High |
| 499333395 [231-244] | 1728.8949 | 2.08 | 0.00179 | VMKEIENQVPEIQR | High |
| 499333395 [118-126] | 1099.50884 | -2.22 | -0.00122 | VSLSMDYER | High |
| 499333395 [118-126] | 1115.50375 | -2.33 | -0.0013 | VSLSMDYER | High |
| 499333395 [118-127] | 1227.6038 | -2.88 | -0.00177 | VSLSMDYERK | High |
| 499333395 [234-244] | 1354.69612 | -0.56 | -0.00038 | EIENQVPEIQR | High |
| 499333395 [100-112] | 1351.68993 | -1.22 | -0.00082 | HPGANTNMIGLAR | High |
| 499333395 [100-112] | 1367.68484 | -0.34 | -0.00023 | HPGANTNMIGLAR | High |
| 499333395 [293-300] | 1022.6067 | -0.38 | -0.00019 | MKNYILIK | High |
| 499333395 [262-270] | 1177.55242 | -1.66 | -0.00098 | LPYDEFHEK | High |

**~28 kDa band**

**ABC transporter, ATP-binding protein, MA_3998, MW: 59.6**

**Sequence coverage: 46%**

| Positions in Master Proteins | Theo. MH+ [Da] | DeltaM [ppm]: Mascot | Deltam/z [Da]: Mascot | Sequence | Confidence |
|---|---|---|---|---|---|
| 19918090 [415-424] | 1064.5371 | -1.67 | -0.00089 | AAGFEENKAK | High |
| 19918090 [470-478] | 944.57751 | -0.78 | -0.00037 | VAVTNSILK | High |
| 19918090 [44-51] | 998.58155 | 0.17 | 0.00008 | TILMHVLR | High |
| 19918090 [124-132] | 1071.51055 | 1.02 | 0.00055 | TFALYGDER | High |
| 19918090 [39-51] | 1414.78349 | 0.92 | 0.00043 | SGSGKTILMHVLR | High |
| 19918090 [192-211] | 2126.10495 | 0.96 | 0.00102 | NPLLLLADEPTGTLDPMTAK | High |
| 19918090 [9-18] | 1121.58372 | -1.24 | -0.0007 | NLTVDFDGLK | High |
| 19918090 [430-441] | 1509.60567 | 1.1 | 0.00056 | MTDEMSEGERHR | High |
| 19918090 [454-469] | 1760.88088 | 1.05 | 0.00093 | IVIMDEPTGTMDPITK | High |
| 19918090 [212-222] | 1206.72048 | 3.36 | 0.00203 | LVHEAIINAVK | High |
| 19918090 [103-111] | 1104.64117 | 1.32 | 0.00073 | LSLYDPIRK | High |
| 19918090 [27-38] | 1227.69433 | -1.15 | -0.0007 | INEGEVVGILGK | High |
| 19918090 [361-368] | 858.47919 | 0.19 | 0.00008 | LGVDNKGR | High |
| 19918090 [117-123] | 860.50223 | 0.08 | 0.00004 | IAIMLQR | High |
| 19918090 [442-453] | 1350.84675 | -3 | -0.00202 | IALAQVLIKEPR | High |
| 19918090 [154-167] | 1636.9017 | -1.44 | -0.00118 | KAVELLEEVNLSHR | High |
| 19918090 [52-69] | 2021.99269 | -1.92 | -0.0013 | GTESFENISGEVIYHLAR | High |
| 19918090 [174-182] | 1003.5167 | -0.35 | -0.00018 | ELSGGEKQR | High |

| | | | | | |
|---|---|---|---|---|---|
| 19918090 [315-329] | 1433.76347 | -1.44 | -0.00104 | EGEIFGLVGISGAGK | High |
| 19918090 [73-81] | 1109.5408 | 0.89 | 0.0005 | CGYIERPSK | High |
| 19918090 [155-167] | 1508.80673 | -1.89 | -0.00143 | AVELLEEVNLSHR | High |
| 19918090 [335-350] | 1726.94078 | -1.8 | -0.00156 | ILMGILPPTSGEVEVR | High |
| 19918090 [295-304] | 1163.65313 | 0.1 | 0.00006 | YISVDRGVVR | High |

**~25 kDa band**

**ABC transporter, ATP-binding protein, MA_3998, MW: 59.6**

**Sequence coverage: 16%**

| Positions in Master Proteins | Theo. MH+ [Da] | DeltaM [ppm]: Mascot | Deltam/z [Da]: Mascot | Sequence | Confidence |
|---|---|---|---|---|---|
| 19918090 [415-424] | 1064.5371 | 0.05 | 0.00003 | AAGFEENKAK | High |
| 19918090 [27-38] | 1227.69433 | 0.64 | 0.00039 | INEGEVVGILGK | High |
| 19918090 [361-368] | 858.47919 | -0.16 | -0.00007 | LGVDNKGR | High |
| 19918090 [442-453] | 1350.84675 | -1.82 | -0.00123 | IALAQVLIKEPR | High |
| 19918090 [174-182] | 1003.5167 | 0.26 | 0.00013 | ELSGGEKQR | High |
| 19918090 [73-81] | 1109.5408 | -2.3 | -0.00127 | CGYIERPSK | High |
| 19918090 [19-26] | 927.57343 | 0.77 | 0.00036 | ALKNVNLR | High |
| 19918090 [454-469] | 1760.88088 | 0.98 | 0.00087 | IVIMDEPTGTMDPITK | High |

**Methanogenesis Marker Protein 17, MA_3993, MW: 22.7**

**Sequence coverage: 19%**

| Positions in Master Proteins | Theo. MH+ [Da] | DeltaM [ppm]: Mascot | Deltam/z [Da]: Mascot | Sequence | Confidence |
|---|---|---|---|---|---|
| 499333396 [23-32] | 1115.63067 | -0.99 | -0.00055 | IIEDNLASLK | High |
| 499333396 [141-151] | 1340.73211 | 1 | 0.00067 | LHENLVEFAIR | High |

| | | | | | |
|---|---|---|---|---|---|
| 499333396 [33-39] | 697.43554 | 0 | 0 | LAPAIGR | High |
| 499333396 [22-32] | 1243.72563 | 0 | 0 | KIIEDNLASLK | High |
| 499333396 [152-160] | 1032.5585 | -1.52 | -0.00079 | ATPEGFRVR | High |

**~16 kDa band**

| **conserved hypothetical protein, MA_4200, MW: 17.9** | | | | | |
|---|---|---|---|---|---|
| **Sequence coverage: 64%** | | | | | |
| Positions in Master Proteins | Theo. MH+ [Da] | DeltaM [ppm]: Mascot | Deltam/z [Da]: Mascot | Sequence | Confidence |
| 19918309 [44-51] | 946.53564 | -0.4 | -0.00019 | AFKELNPK | High |
| 19918309 [1-17] | 1884.00076 | 3.77 | 0.00355 | MIGIISDSHDNLTAIRK | High |
| 19918309 [1-16] | 1771.90071 | -0.08 | -0.00007 | MIGIISDSHDNLTAIR | High |
| 19918309 [1-16] | 1755.90579 | 1.82 | 0.00159 | MIGIISDSHDNLTAIR | High |
| 19918309 [126-145] | 2046.98345 | -3.22 | -0.0033 | MIDGTPVINPGECSGVLSGK | High |
| 19918309 [126-145] | 2030.98854 | -1.18 | -0.0012 | MIDGTPVINPGECSGVLSGK | High |
| 19918309 [52-68] | 1930.9909 | -1.15 | -0.00074 | LYFVFGNNDGDKVTLTK | High |
| 19918309 [1-17] | 1899.99567 | 0.02 | 0.00001 | MIGIISDSHDNLTAIRK | High |
| 19918309 [52-63] | 1388.64811 | -1.05 | -0.00073 | LYFVFGNNDGDK | High |
| 19918309 [117-125] | 975.4755 | 0.09 | 0.00003 | GHTHDPGVR | High |
| 19918309 [29-43] | 1595.8904 | -1.74 | -0.00139 | AVLHAGDIISPFTVR | High |
| 19918309 [18-25] | 982.53564 | 0.48 | 0.00024 | AVEFFNKK | High |
| 19918309 [18-24] | 854.44068 | 0.62 | 0.00026 | AVEFFNK | High |

| | | | | | |
|---|---|---|---|---|---|
| 19918309 [104-116] | 1377.70087 | -1.34 | -0.00092 | ALAESGDFDVVVR | High |

**Table S3.** Identification of individual SDS-PAGE bands of in-gel digests from MCR preparation showing tryptic peptides and resulting proteins identified.

**~60 kDa band**

| **ABC transporter, ATP-binding protein, MA_3998, MW: 59.6** | | | | | |
|---|---|---|---|---|---|
| **seq coverage: 65.9%** | | | | | |
| Conf % | Observ. mass | Calc. mass | Ion% | # | Sequence |
| 100.00% | 963.71686 | 962.53796 | 76.90% | 2 | -.MPVFIEVK.N |
| 18.20% | 979.93097 | 978.53284 | 75.00% | 1 | -.M(15.9949)PVFIEVK.N |
| 100.00% | 1121.976 | 1121.5837 | 100.00% | 7 | K.NLTVDFDGLK.A |
| 100.00% | 1228.6326 | 1227.6943 | 100.00% | 5 | R.INEGEVVGILGK.S |
| 99.80% | 983.73737 | 982.5866 | 92.30% | 14 | K.TILMHVLR.G |
| 57.80% | 999.5153 | 998.5815 | 84.60% | 2 | K.TILM(15.9949)HVLR.G |
| 100.00% | 2023.1487 | 2021.9927 | 57.10% | 3 | R.GTESFENISGEVIYHLAR.C |
| 100.00% | 2023.6553 | 2021.9927 | 57.40% | 16 | R.GTESFENISGEVIYHLAR.C |
| 100.00% | 1931.0596 | 1929.8608 | 57.70% | 2 | K.CPVCGETLEEFEADFIK.L |
| 59.20% | 1929.4131 | 1929.8608 | 45.50% | 1 | K.CPVCGETLEEFEADFIK.L |
| 99.70% | 977.57355 | 976.5462 | 100.00% | 2 | K.LSLYDPIR.K |
| 98.40% | 1105.427 | 1104.6411 | 46.20% | 17 | K.LSLYDPIRK.D |
| 11.90% | 1106.187 | 1104.6411 | 50.00% | 1 | K.LSLYDPIRK.D |
| 99.70% | 845.13745 | 844.5073 | 100.00% | 8 | R.IAIMLQR.T |

| | | | | | |
|---|---|---|---|---|---|
| 76.60% | 861.46533 | 860.5022 | 100.00% | 4 | R.IAIM(15.9949)LQR.T |
| 100.00% | 1071.9497 | 1071.5105 | 100.00% | 5 | R.TFALYGDER.V |
| 100.00% | 1675.6926 | 1674.9425 | 87.50% | 9 | R.VLVNVINSLNEIGYK.G |
| 100.00% | 1676.0977 | 1674.9425 | 72.50% | 2 | R.VLVNVINSLNEIGYK.G |
| 99.60% | 2266.596 | 2268.1653 | 36.50% | 3 | K.GEDAMKKAVELLEEVNLSHR.M |
| 28.40% | 1637.2201 | 1636.9016 | 45.90% | 1 | K.KAVELLEEVNLSHR.M |
| 100.00% | 1509.824 | 1508.8068 | 95.00% | 31 | K.AVELLEEVNLSHR.M |
| 25.90% | 1508.7313 | 1508.8068 | 48.50% | 1 | K.AVELLEEVNLSHR.M |
| 29.10% | 2264.4473 | 2266.143 | 35.40% | 2 | K.AVELLEEVNLSHRM(15.9949)M(15.9949)HVAR.E |
| 100.00% | 2110.84 | 2110.11 | 72.70% | 2 | R.NPLLLLADEPTGTLDPMTAK.L |
| 100.00% | 2112.1482 | 2110.11 | 61.10% | 2 | R.NPLLLLADEPTGTLDPMTAK.L |
| 100.00% | 2127.9412 | 2126.105 | 81.80% | 4 | R.NPLLLLADEPTGTLDPM(15.9949)TAK.L |
| 100.00% | 2127.4265 | 2126.105 | 56.40% | 2 | R.NPLLLLADEPTGTLDPM(15.9949)TAK.L |
| 100.00% | 1207.6759 | 1206.7205 | 94.10% | 36 | K.LVHEAIINAVK.N |
| 20.60% | 2179.9954 | 2178.036 | 28.30% | 1 | K.NYNMSMVLTSHWPEVIEK.L |
| 100.00% | 3262.604 | 3260.654 | 33.80% | 2 | K.AILLENGEVVQEGDPLEVSAIFMQSVSMVR.Q |
| 97.00% | 3278.1309 | 3276.649 | 28.40% | 1 | K.AILLENGEVVQEGDPLEVSAIFM(15.9949)QSVSMVR.Q |
| 100.00% | 3277.5852 | 3276.649 | 41.60% | 2 | K.AILLENGEVVQEGDPLEVSAIFMQSVSM(15.9949)VR.Q |
| 100.00% | 3294.1882 | 3292.6438 | 30.70% | 2 | K.AILLENGEVVQEGDPLEVSAIFM(15.9949)QSVSM(15.9949)VR.Q |
| 100.00% | 1110.2744 | 1109.6677 | 100.00% | 9 | K.NVLVGEPIIR.V |

| | | | | | |
|---|---|---|---|---|---|
| 92.20% | 2177.3682 | 2176.2197 | 34.60% | 1 | R.DLSKRYISVDRGVVRAVDK.I |
| 40.40% | 909.31757 | 908.4948 | 75.00% | 1 | K.RYISVDR.G |
| 100.00% | 1434.0156 | 1433.7634 | 100.00% | 3 | K.EGEIFGLVGISGAGK.T |
| 100.00% | 1435.8513 | 1433.7634 | 56.10% | 2 | K.EGEIFGLVGISGAGK.T |
| 11.20% | 1954.9984 | 1952.0334 | 29.60% | 1 | K.EGEIFGLVGISGAGKTTTSK.I |
| 100.00% | 1711.7412 | 1710.9458 | 100.00% | 6 | K.ILMGILPPTSGEVEVR.V |
| 30.70% | 1713.7292 | 1710.9458 | 31.80% | 1 | K.ILMGILPPTSGEVEVR.V |
| 100.00% | 1727.6694 | 1726.9408 | 84.60% | 7 | K.ILM(15.9949)GILPPTSGEVEVR.V |
| 100.00% | 1180.3419 | 1179.535 | 86.70% | 2 | R.VGDEWVDMTK.L |
| 90.70% | 657.8392 | 658.44977 | 88.90% | 2 | K.AIITLK.A |
| 43.30% | 1264.2993 | 1261.8103 | 50.00% | 1 | R.HRIALAQVLIK.E |
| 87.00% | 970.3972 | 968.65027 | 100.00% | 4 | R.IALAQVLIK.E |
| 100.00% | 1761.7612 | 1760.8809 | 92.00% | 4 | R.IVIMDEPTGTMDPITK.V |
| 100.00% | 1777.364 | 1776.8757 | 96.00% | 4 | R.IVIMDEPTGTM(15.9949)DPITK.V |
| 100.00% | 1777.4922 | 1776.8757 | 72.00% | 4 | R.IVIM(15.9949)DEPTGTMDPITK.V |
| 99.70% | 947.56476 | 944.5775 | 93.30% | 4 | K.VAVTNSILK.A |
| 41.00% | 688.7841 | 686.41956 | 80.00% | 1 | K.IVEIGR.P |
| 100.00% | 2111.9724 | 2110.1501 | 58.50% | 7 | K.IVEIGRPGTVLAQLTEEER.I |
| 100.00% | 1443.5706 | 1442.7485 | 95.00% | 2 | R.PGTVLAQLTEEER.I |

**Table S4** A list of all primers and gblocks used in this study

| Name | Sequence |
|---|---|
| SA025 | TCATGGTCACATCCTCAGTTTGAAAAAGGTGGAGGACCAGTATTTATTGAAGTTAAAAA |
| SA026 | TGAAAAAGGTGGCGGGTCAGGCGGGGGTTCAGGGGGTGGCTCATGGTCACATCCTCAG |
| SA027 | ACGACAAGGGTTCCGCTGCATCCTGGTCACATCCTCAGTTTGAAAAAGGTGGCGGG |
| SA028 | AATAAATTAAGGAGGAAATTCAATGGACTACAAAGACGATGACGACAAGGGTTCCGC |
| SA029 | CATTATACGAAGTTATCAAGATTATATTTCTTCTGCAGCTTTGATCCT |
| SA030 | AATAAATTAAGGAGGAAATTCAATGCCAGTATTTATTGAAGTT |
| SA031 | GCGGAACCCTTGTCGTCATCGTCTTTGTAGTCTCCTCCACCTATTTCTTCTGCAGCTTTG |
| SA032 | CTGACCCGCCACCTTTTTCAAACTGAGGATGTGACCAGGATGCAGCGGAACCCTTGTCGT |
| SA033 | TCAAACTGAGGATGTGACCATGAGCCACCCCCTGAACCCCCGCCTGACCCGCCACCTTT |
| SA034 | CATTATACGAAGTTATCAAGATTATTTTTCAAACTGAGGATGTGACCA |
| SA119 | CTTGACCCCATGACTGCAAA |
| SA120 | AATTTTCTCCAGATCCGAGACTTT |
| SA124 | AGACCGGGATTCCATTTCCT |
| SA126 | TATTTGCAAGCTCGGAAAGGA |
| SA131 | TCTCGGTAGAAGACGGAATTCC |
| SA134 | CTGGAAACCCCGGCTTCTT |
| SA137 | TGCAGGGGAAATCGGGATTA |
| SA138 | CAGGACTTGCCCAGTGGATA |
| SA148 | GCATCAGGATTGTGGCCCCTGATCCGCTTTT |
| SA149 | GCGGATCAGGGGCCACAATCCTGATGCAC |
| SA150 | AATAAATTAAGGAGGAAATTCA |
| SA151 | AGGTTGTGGTGGCGCCTGCTCCGCTTAT |
| SA152 | AAGCGGAGCAGGCGCCACCACAACCTCTAAAAT |
| SA173 | CTTCTGTTAGCTGACGCGCCTACGGGTACA |
| SA174 | TGTACCCGTAGGCGCGTCAGCTAACAGAAG |
| SA175 | GTTATTATGGACGCACCTACAGGGACCATGGAC |
| SA176 | GTCCATGGTCCCTGTAGGTGCGTCCATAATAAC |

| | |
|---|---|
| gSAA001 | AATAAATTAAGGAGGAAATTCAATGGACTACAAAGACGATGACGACA<br>AGGGTTCCGCTGCATCCTGGTCACATCCTCAGTTTGAAAAAGGTGGCG<br>GGTCAGGCGGGGGTTCAGGGGGTGGCTCATGGTCACATCCTCAGTTT<br>GAAAAAGGTGGAGGACCAGTATTTATTGAAGTTAAAAACCTAACTGTA<br>GACTTTGACGGTCTCAAAGCCCTGAAAAATGTAAATTTAAGGATCAAC<br>GAAGGGGAAGTCGTTGGAATTCTGGGAAAAAGCGGATCAGGGAAAA<br>CAATCCTGATGCACGTGTTGCGCGGGACCGAATCTTTTGAGAATATCTC<br>AGGCGAAGTAATCTATCATCTGGCCCGCGCAGAGAAAGCTGGATATAT<br>AGAGCGTCCAAGCAAAATAGGCCAGAAAGCCCCGGTCGCAGGGGAG<br>ACGCTTGAGGAGTTCGAAGCTG |

**Table S5** List of all plasmids used in this study.

| Plasmid | Description | Reference |
|---|---|---|
| pJK027A | Vector with PmcrB(tetO1) promoter fusion to *uidA* that contains φC31-attB and λattP | (3) |
| pAMG40 | Vector for fosmid retrofitting containing pC2A and λattB | (3) |
| pDN201 | pJK027A-derived plasmid with P*mcrB*(tetO1) promoter fusion to Spy Cas9 | (4) |
| pSAA004 | derivative of pJK027A with MA3998_N_1XFLAG_2XStrep under control of P*mcrB*(tetO1) | This study |
| pSAA049 | derivative of pJK027A with MA3998_C_1XFLAG_2XStrep under control of P*mcrB*(tetO1) | This study |
| pSAA050 | derivative of pJK027A with MA3998_N_1XFLAG_2XStrep_K43A under control of P*mcrB*(tetO1) | This study |
| pSAA051 | derivative of pJK027A with MA3998_N_1XFLAG_2XStrep_K329A under control of P*mcrB*(tetO1) | This study |
| pSAA052 | derivative of pJK027A with MA3998_N_1XFLAG_2XStrep_K43/329A under control of P*mcrB*(tetO1) | This study |
| pSAA055 | derivative of pJK027A with MA3998_N_1XFLAG_2XStrep_E200A under control of P*mcrB*(tetO1) | This study |
| pSAA056 | derivative of pJK027A with MA3998_N_1XFLAG_2XStrep_E459A under control of P*mcrB*(tetO1) | This study |
| pSAA057 | derivative of pJK027A with MA3998_N_1XFLAG_2XStrep_E200/459A under control of P*mcrB*(tetO1) | This study |
| pSAA058 | derivative of pJK027A with MA3998_N_1XFLAG_2XStrep_C70/73/86/89A under control of P*mcrB*(tetO1) | This study |
| pDN353 | pDN201 derived plasmid containing the sgRNA and repair template to generate an in-frame deletion of MA_3992 | This study |
| pDN355 | pDN201 derived plasmid containing the sgRNA and repair template to generate an in-frame deletion of MA_3993 | This study |
| pDN356 | pDN201 derived plasmid containing the sgRNA and repair template to generate an in-frame deletion of MA_3994 | This study |
| pDN357 | pDN201 derived plasmid containing the sgRNA and repair template to generate an in-frame deletion of MA_3995 | This study |
| pDN358 | pDN201 derived plasmid containing the sgRNA and repair template to generate an in-frame deletion of MA_3996 | This study |
| pDN359 | pDN201 derived plasmid containing the sgRNA and repair template to generate an in-frame deletion of MA_3997 | This study |

| | | |
|---|---|---|
| pDN360 | pDN201 derived plasmid containing the sgRNA and repair template to generate an in-frame deletion of MA_3998 | This study |
| pDN367 | Cointegrate of pDN353 and pAMG40 obtained using Gateway cloning (BP Clonase II) | This study |
| pDN369 | Cointegrate of pDN355 and pAMG40 obtained using Gateway cloning (BP Clonase II) | This study |
| pDN371 | Cointegrate of pDN357 and pAMG40 obtained using Gateway cloning (BP Clonase II) | This study |
| pDN373 | Cointegrate of pDN359 and pAMG40 obtained using Gateway cloning (BP Clonase II) | This study |
| pDN375 | Cointegrate of pDN361 and pAMG40 obtained using Gateway cloning (BP Clonase II) | This study |
| pDN377 | Cointegrate of pDN363 and pAMG40 obtained using Gateway cloning (BP Clonase II) | This study |
| pDN379 | Cointegrate of pDN365 and pAMG40 obtained using Gateway cloning (BP Clonase II) | This study |

**Table S6** List of all the strains used in this study

| Strain | Description | Reference |
|---|---|---|
| WWM60 | *M. acetivorans* Δ*hpt*:: P*mcrB-tetR* | (3) |
| WWM1086 | *M. acetivorans* Δ*hpt*:: P*mcrB-tetR* TAP (3X-FLAG EK site 2X Strep EK site) tag at N-terminus of *mcrG* (WWM60/pDN329) | (5) |
| WWM73 | *M. acetivorans* Δ*hpt*:: P*mcrB-tetR* -phiC31int-attP | (3) |
| DDN122 | *M. acetivorans* Δ*hpt*:: P*mcrB-tetR*-phiC31int-attP att::P*mcrB*(tetO1) NstrepFLAG- MA3998 (WWM73/att:pSAA004) | this study |
| DDN291 | *M. acetivorans* Δ*hpt*:: P*mcrB-tetR* -phiC31int-attP att::P*mcrB*(tetO1) MA3998-CstrepFLAG (WWM73/ att:pSAA049) | this study |
| DDN324 | M. acetivorans delta hpt:: PmcrB-tetR-phiC31int-attP att::PmcrB(tetO1) MA3998-NstrepFLAG-K43A (WWM73/ att:pSAA050) | this study |
| DDN325 | *M. acetivorans* Δ*hpt*:: P*mcrB-tetR* -phiC31int-attP att::P*mcrB*(tetO1) MA3998-NstrepFLAG-K329A (WWM73/ att:pSAA051) | this study |
| DDN326 | *M. acetivorans* Δ*hpt*:: P*mcrB-tetR* -phiC31int-attP att::P*mcrB*(tetO1) MA3998-NstrepFLAG-K43A-K329A (WWM73/ att:pSAA052) | this study |
| DDN408 | *M. acetivorans* Δ*hpt*:: P*mcrB-tetR* -phiC31int-attP att::P*mcrB*(tetO1) MA3998-NstrepFLAG-E200A (WWM73/ att:pSAA055) | this study |
| DDN409 | *M. acetivorans* Δ*hpt*:: P*mcrB-tetR* -phiC31int-attP att::P*mcrB*(tetO1) MA3998-NstrepFLAG-E459A (WWM73/ att:pSAA056) | this study |
| DDN410 | *M. acetivorans* Δ*hpt*:: P*mcrB-tetR* -phiC31int-attP att::P*mcrB*(tetO1) MA3998-NstrepFLAG-E200/459A (WWM73/ att:pSAA057) | this study |
| DDN435 | *M. acetivorans* Δ*hpt*:: P*mcrB-tetR* -phiC31int-attP att::P*mcrB*(tetO1) MA3998-NstrepFLAG-C70/73/86/89 (WWM73/ att:pSAA058) | this study |

Supplemental References

1. G. L. Chadwick, G. A. Dury, D. D. Nayak, Physiological and transcriptomic response to methyl-coenzyme M reductase limitation in Methanosarcina acetivorans. *Applied and Environmental Microbiology* **90**, e0222023 (2024).

2. K. E. Shalvarjian, *et al.*, Methanogenic archaea encoding Pyrrolysine maintain ambiguous amber codon usage. *Proceedings of the National Academy of Sciences* **122**, e2517473122 (2025).

3. A. M. Guss, M. Rother, J. K. Zhang, G. Kulkkarni, W. W. Metcalf, New methods for tightly regulated gene expression and highly efficient chromosomal integration of cloned genes for *Methanosarcina* species. *Archaea* **2**, 193–203 (2008).

4. D. D. Nayak, W. W. Metcalf, Cas9-mediated genome editing in the methanogenic archaeon Methanosarcina acetivorans. *Proceedings of the National Academy of Sciences of the United States of America* **114**, 2976–2981 (2017).

5. D. D. Nayak, N. Mahanta, D. A. Mitchell, W. W. Metcalf, Post-translational thioamidation of methyl-coenzyme M reductase, a key enzyme in methanogenic and methanotrophic Archaea. *eLife* **6**, e29218 (2017).
